# Multi-modal screening for synergistic neuroprotection of extremely preterm brain injury

**DOI:** 10.1101/2025.04.25.650638

**Authors:** Zheyu Ruby Jin, Kylie A. Corry, Olivia C. Brandon, Matthew J. Magoon, Hawley Helmbrecht, Daniel H. Moralejo, Robell Bassett, Sarah E. Kolnik, Patrick M. Boyle, Sandra E. Juul, Elizabeth Nance, Thomas R. Wood

## Abstract

Preterm brain injury affects both white and grey matter, including altered cortical development and gyrification, with associated neurodevelopmental sequelae such as cerebral palsy and learning deficits. The preterm brain also displays regionally heterogeneous responses to both injury and treatment, supporting the need for drug combinations to provide global neuroprotection. We developed an extremely preterm-equivalent organotypic whole hemisphere (OWH) slice culture injury model using the gyrencephalic ferret brain to probe treatment mechanisms of promising therapeutic agents and their combination. Regional and global responses to injury and treatment were assessed by cell death quantification, machine learning-augmented morphological microglia assessments, and digital transcriptomics. Using two promising therapeutic agents, azithromycin (Az) and erythropoietin (Epo), we show minimal neuroprotection by either therapy alone, but evidence of synergistic neuroprotection by Az*Epo both globally and regionally. This effect of Az*Epo involved emergent augmentation of transcriptomic responses to injury related to neurogenesis and neuroplasticity and downregulation of transcripts involved in cytokine production, inflammation, and cell death. This study supports the use of the ferret OWH slice culture model to provide a powerful high-throughput platform to examine combinations of therapeutics for extremely preterm brain injury.

## Introduction

Preterm birth is one of the leading causes of neonatal morbidity and mortality worldwide (*1*), with more than 10% of infants born preterm globally. Perinatal infection, hypoxia-ischaemia (HI), and hyperoxia are common during and after preterm birth, contributing to inflammatory and oxidative brain injury (*2*). Extremely preterm infants, born before 28 weeks’ gestation, experience a high rate of death and disability; at least 10% die and >60% of the survivors develop at least one disability such as cerebral palsy, autism, attention-deficit/hyperactivity disorder (ADHD), or cognitive, hearing, or visual impairment (*3–6*). Individuals born extremely preterm are also at a higher risk of several chronic diseases including mental health disorders and cognitive impairment, thought to be due to alterations in immune, metabolic, and brain function resulting from exposures in the perinatal and neonatal period (*7, 8*). However, there are currently no targeted neuroprotective interventions for preterm infants, and as preterm survival rate continues to improve (*9*), it is critical to develop therapies to optimise neurodevelopmental outcomes for this population.

Preterm brain injury is characterised by both white matter injury (WMI) and grey matter injury (*10, 11*), including altered cortical development and gyrification (*12*). Due to the nature of the brain injury seen in preterm infants, the ferret is a highly relevant animal model in which to study preterm brain injury. Unlike rodents and rabbits, ferrets are born lissencephalic and develop a gyrencephalic cerebral cortex postnatally (*13*). The ferret brain undergoes postnatal white matter maturation and complex cortical folding, a process that is not present in rodents, in a similar pattern as the human brain during the third trimester (*14*). We have previously shown that an *in vivo* ferret model of preterm brain injury (*13*) displays WMI, altered gyrification, and behavioural changes consistent with those seen in premature infants (*8, 15*).

Due to the urgent need for neuroprotective therapies for extremely preterm infants as well as the ever-increasing list of potential therapeutic candidates, high-throughput methods to test drugs and drug combinations are needed. Organotypic whole-hemisphere (OWH) slice models provide a platform to bridge the gap between *in vitro* cell lines and *in vivo* models by maintaining the brain’s structural integrity and capturing regional variability in susceptibility to injury and therapeutic responses (*16*). Prior work in a term-equivalent ferret OWH slice model of oxygen-glucose deprivation (OGD) to simulate ischemic neuronal injury showed regional dependence in response to injury and treatment, while mirroring trends seen after injury *in vivo* (*17*). The OWH slice model is also amendable to studying glial cell dynamics critical for neuroplasticity regulation (*18*), as well as exploration of global and regional responses to injury such as transcriptomics (*19, 20*). Therefore, the ferret OWH slice culture platform provides a promising method to screen and compare therapies for preterm brain injury at scale before *in vivo* assessment.

In this study, we developed an extremely preterm-equivalent ferret OWH brain slice model of OGD to investigate injury responses and the neuroprotective effects of promising candidates for neuroprotection. As preterm brain injury patterns are heterogeneous and display regional responses to therapy, it is increasingly acknowledged that drug combinations - rather than single therapies – are likely to be needed to provide robust global neuroprotection. We use azithromycin (Az) (*21*) and erythropoietin (Epo) (*22*) and their combination (Az*Epo) to perform global and regional cellular analyses and targeted transcriptomics. Az and Epo are already used in preterm infants for anaemia of prematurity and treatment of ureaplasma infections, respectively. Epo is neuroprotective and neuroregenerative in animal models of neonatal brain injury (*23, 24*). Although results from the recent Preterm Epo Neuroprotection Trial (PENUT) showed no significant difference in death or neurodevelopmental impairment between placebo and Epo-treated groups of extremely preterm infants (*25*), Epo remains a promising candidate to be used in combination with other therapies (*26–28*). Az reduces injury severity in adult rodent stroke and spinal cord injury models as well as neonatal HI models (*21, 29*), contributing to early neuroprotection and long-term amelioration of morbidity by modulating microglial and macrophage phenotype (*30, 31*). As WMI in the preterm brain is thought to be associated with the location of dense microglial aggregates as they migrate through the central nervous system during brain development (*32, 33*), modulation of microglial phenotype with Az is a promising potential intervention for the injured extremely preterm brain. Here, we topically apply Az and Epo to OGD-exposed OWH ferret brain slices and measure regional and global cell death, microglial density and morphology, and transcriptomic profiles. Using a combination of robust statistical models and both supervised and unsupervised machine learning techniques, we show that Az*Epo results in emergent synergistic neuroprotection, including transcriptomic and neuroprotective responses that cannot be predicted from the action of each therapy individually. Our results support the use of the ferret OWH model to screen promising treatment combinations for neonatal brain injuries in a high-throughput manner, providing an accelerated pathway for clinical translation.

## Materials and Methods

### Animal care and ethics

This study was performed in strict accordance with the National Institutes of Health Guide for the Care and Use of Laboratory Animals. All animals were handled according to an approved Institutional Animal Care and Use Committee (IACUC) protocol (#3328-06) of the University of Washington. The University of Washington has an approved Animal Welfare Assurance (#A3464-01) on file with the National Institute of Health Office of Laboratory Animal Welfare, is registered with the United States Department of Agriculture (USDA, certificate #91-R-0001), and is accredited by American Association for Accreditation of Laboratory Animal Care (AALAC) International. Ferret jills with cross-fostered kits were purchased from Marshall BioResources (North Rose, NY, USA) and arrived at the facility at or before postnatal day (P) 8. Animals were maintained in a centralised vivarium and had *ad libitum* access to food and water. Standard housing conditions included a 16 h light/8 h dark cycle with a room temperature range of 61–72 °F (16–22 °C), humidity of 30%–70%, and 10–15 fresh air changes per hour.

### Organotypic whole hemisphere (OWH) ferret brain slice preparation

Full experimental procedures are depicted in **Supplemental Figure S1**. At P14, comparable to extremely preterm (<28 weeks’) human gestation, ferret kits were deeply anesthetised using 5% isoflurane and administered an overdose intraperitoneal injection of pentobarbital (120–150 mg/kg). Animals were then quickly decapitated using a guillotine, and whole brains were extracted, cut into hemispheres, and placed into ice-cold dissecting media consisting of 0.64% w/v glucose, 100% Hank’s Balanced Salt Solution (HBSS), and 1% penicillin–streptomycin. Live whole-hemisphere 300 μm slices were obtained using a Leica Vibratome. Slices were immediately transferred to 35 mm, 0.4 μm-pore membrane inserts in six-well plates and cultured in 1 ml of 5% heat-inactivated horse-serum slice culture media (SCM) consisting of 50% Minimum Essential Media (MEM), 45% HBSS, 1% GlutaMAX, and 1% penicillin–streptomycin. Slices in the normal control (NC) group were not subjected to oxygen-glucose deprivation (OGD) and were maintained in SCM with 5% heat-inactivated horse serum for the duration of the study. Slices were cultured in a sterile incubator at constant temperature (37°C), humidity, and CO_2_ level (5%).

### Oxygen-glucose deprivation

After 3 days *in vitro* (DIV), all non-NC slices were subjected to 2 h OGD injury, adapted from an *ex vivo* rat brain slice OGD platform, resulting in measurable injury without complete cell death (*34*). SCM was replaced with glucose-free OGD media containing 150 mM NaCl, 2.8 mM KCl, 1 mM CaCl_2_, and 10 mM HEPES (4-(2-hydroxyethyl)-1piperazineethanesulfonic acid (HEPES) in DI H_2_O. OGD media was first prewarmed to 37°C and bubbled for 5 min with N_2_ at a flow rate of 3 L/min to deprive it of O_2_. The slices were then transferred into a sealable chamber. The chamber was flushed with N_2_ for 10 min at a flow rate of 5 L/min before being placed back into the 37°C sterile incubator for 2 h. Slices were then removed from the hypoxic chamber and the media was replaced with 5% SCM or SCM containing 15 μM Az or/and 3 IU/ml Epo, which are concentrations previously shown to be neuroprotective *in vitro*. Treatment-containing media was added 100 μl topically onto the slices, and 900 μl under the inserts. For all studies, the end of OGD incubation was defined as time t = 0 h. The slices were cultured for an additional 24 h under normoxic conditions (5% CO_2_, balance air). 5% SCM was replaced for the NC slices at the same time points as non-NC slices to match the number of media changes. Supernatant was collected during media changes at the end of OGD (t = 0 h) and at the end of culturing (t = 24 h) to perform cell death assays and immunofluorescent imaging.

### Lactate dehydrogenase (LDH) assay

Supernatant collected at t = 0 h and 24 h was immediately placed at -80°C. Prior to running the LDH assay (Cayman Chemical, Ann Arbor, MI, USA), the supernatant was thawed at room temperature. LDH is an enzyme released from cells upon membrane degradation in response to cytotoxicity. Through a series of coupled enzymatic reactions, LDH can be converted to formazan, which absorbs in the 490–520 nm range. 100 μl of supernatant from each slice was seeded in triplicate into a 96-well plate, followed by addition of 100 μl chilled LDH reaction buffer to each well. The plate was incubated at 37°C for 30 min. Absorbance was measured at 490 nm (A_490_) on the UV–Vis Spectrophotometer. A_490_ for t = 24 h was reported to give a measure of post-treatment LDH release. Supernatant was tested for n = 12 slices per group with an equal sex split.

### Cell viability quantification

At t = 24 h, slices were fixed in 10% buffered formalin and then stored in 1 × phosphate-buffered saline (PBS) at 4°C. The slices were stained with 1 mL of 5 μg/mL 4’,6-diamidino-2-phenylindole (DAPI, Invitrogen) in 1 × PBS for 15 min then washed twice in 1 ×PBS. Using a Nikon A1R confocal microscope with the 40× objective, representative images of the stained slices were obtained in four different regions, comprising the cortex, subcortical white matter, thalamus, and basal ganglia. For each experimental group, n = 9-12 slices were imaged with 3 images per region of interest. Cells undergo chromatin and nucleus condensation during apoptosis and necrosis. Since DAPI stain is associated with double-stranded DNA structures, pyknotic nuclei occur from preprogrammed cell death as spheres of compact and dark-staining nuclear chromatin. Cell death was quantified with pyknotic nuclei counts identified in the DAPI-stained regions of interest (ROI).

#### Automated Nucleus Counts

DAPI-stained images were analyzed using a custom computational utility (coded using Python v3.10.11) to automatically calculate total and pyknotic nucleus counts. Briefly, two-dimensional histology images (512x512 pixels, each 0.863x0.863 µm^2^) were preprocessed by normalizing pixel intensities with respect to the image and then applying a high-pass Gaussian spatial filter parameterized with sigma = 1 pixel (SciPy).(*35*) Pixels with a relative intensity <10% were masked and nuclei were detected with the Laplacian of Gaussian method (scikit-image),(*36*) parameterized with the *threshold* at 0.5% of the maximum pixel intensity and the other parameters as follows: *min_sigma* = 0.863, *max_sigma* = 5.179, *num_sigma* = 25, *overlap* = 0.7, *threshold_rel* = 0. Watershed (scikit-image) segmentation further delineated boundaries between clusters of nuclei.(*36*) Eight features were calculated to describe each identified nucleus: area (*A*, Eqn. 1), perimeter (*P*) from the Crofton four-direction approximation (scikit-image), ideal radius (*r_I_*, Eqn. 2), eccentricity (ε, Eqn. 3), average normalized intensity (*I_M_*, Eqn. 4), total normalized intensity (*I_T_*, Eqn. 5), weighted intensity (*I_W_*, Eqn. 6), and the Laplacian of Gaussian sigma parameter value (*σ*, scikit-image) for cells represented by *n* pixels of pixel area *a* = 0.86^2^ µm^2^ each located distance *d* from the cell center.(*36*)

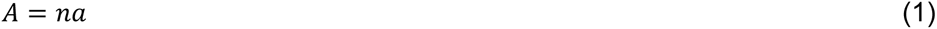

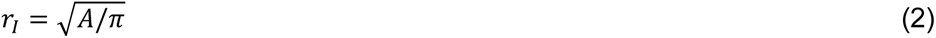

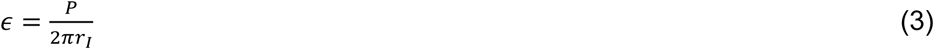

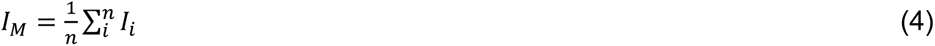

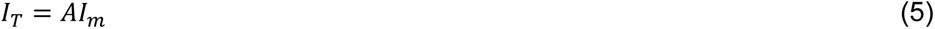

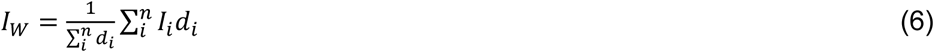

A random forest classifier (scikit-learn)(*37*) was trained on 200 manually annotated DAPI-stained ferret brain histology images that were not used in this study to classify nuclei as pyknotic or non-pyknotic from the eight above-described parameters. Nuclei were identified and characterized by the above methods. To reduce class imbalance, a subset of the data was created with all manually annotated pyknotic nuclei and three times as many randomly selected non-pyknotic nuclei for model development. The classifier was developed from this dataset with 80:20 split 5× cross-validation. It was then used to classify pyknotic nuclei in never-before-seen histology images in this study. The individual creating the code and random forest classifier used in this analysis was fully blinded to both treatment groups and outcomes.

### Immunofluorescent staining

Prior to DAPI staining, the slices were co-stained with a primary antibody for either microglia, rabbit anti-ionised calcium-binding adaptor molecule 1 (rabbit anti-Iba1, Fujifilm) or oligodendrocytes, rabbit anti-oligodendrocyte transcription factor 2, (rabbit anti-Olig2, Sigma-Aldrich) diluted 1:250 in 1×PBS containing 3% Triton X-100 for membrane permeabilization, and 6% goat serum as the blocking reagent. The secondary antibody solution (AF-488 IgG goat anti-rabbit, Invitrogen) was added to the slices at 1:500 in 1×PBS for an additional 2 h at room temperature. Slices were later washed twice again in 1×PBS. Representative images of Iba-1^+^ and Olig2^+^ cells were acquired from n = 6 slices with equal sex split using the Nikon A1R confocal microscope with the 40 × objective. In all experimental groups, 3-5 imaging locations were captured per region including the cortex, subcortical white matter, thalamus, basal ganglia, and corpus callosum.

### Microglial and oligodendrocyte morphological analysis

Images from each ROI of Iba-1-stained slices were converted from .nd2 file format to .tiff file format. Images were then split by colour channel, green for Iba-1 and blue for DAPI. Images from both colour channels were split into four equal quadrants to increase the number of images for training and testing. Converted images underwent a train:test split with a ratio of 80:20 ensuring at least two images from each sex, region, and treatment combination remained after the split. Cells in images were segmented using the Otsu threshold from Sci-kit Image in Python. Objects smaller than 25 pixels were removed, holes were filled, and cells cut off by the edge of the image were removed. Segmented cell images were saved as .png files. The training images were and classification into the machine learning Visual Aided Morpho-Phenotyping Image Recognition (VAMPIRE) pipeline to train a model with a shape mode (SM) number of five and registration coordinates variable set to 50 (*17*). Five SMs were chosen to capture biological variation while remaining computationally efficient. The test images were then run on this model to classify all cells into the five SMs determined during training. Three main morphology parameters of each cell were also determined—perimeter, circularity, and area coverage, defined as the number of pixels within the segmented cell outline. Total number of Iba-1-stained cells per brain structure ROI per slice was estimated by multiplying the total cell counts in the 20% of test images by 5.

### NanoString nCounter RNA extraction and targeted digital transcriptomic quantification

mRNA expression levels by region were analysed with a custom ferret-specific 255 probe digital transcriptomics nCounter panel that included genes related to cell differentiation, inflammation, oxidative stress, and brain development (NanoString, Seattle, USA). As the ferret genome is sequenced, we worked with the NanoString bioinformatics team to develop a custom codeset that was a subset of their standard neuroinflammation and glial pathology panels in the mouse and human. 24 h after OGD and treatment, slices were fixed for 1h in 10% buffered formalin and microdissected into three ROIs: cortex, subcortical white matter, and deep grey matter. mRNA samples were extracted using the Qiagen RNeasy Kit for FFPE (formalin fixed paraffin embedded) tissue, according to the manufacturer’s instructions. The mRNA products were then prepared in nCounter sample wells with the NanoString Master Kit and sent for analyses at the NanoString core facility at the Fred Hutchinson Cancer Research Facility.

### RNA expression profile analysis

Gene ontology (GO) enrichment analyses were performed on differentially expressed transcripts with the ShinyGO graphical gene-set enrichment tool using the ferret genome (*38*). We performed enrichment analysis for Gene Ontology biological process with an adjusted p value <0·05 as the cutoff. A Venn Diagram was computed by the intersection of gene lists described and plotted with InteractiVenn tool (*38*). We used Cytoscape (v3·10·2) to visualise the co-expression network (*39*).

### Statistical analysis

Statistical analyses across treatment groups were performed using the untreated OGD group as the reference group. Comparison of global cell death (LDH) across groups was performed using linear regression with robust standard errors. Comparison of pyknotic nuclei and Iba-1- and Olig1-positive cell counts across groups was performed using linear mixed models with fixed effects of region (global and deep grey matter analyses only) and random effects of slice to account for repeated measures within a given slice/region. Cell count data from the three deep grey regions (thalamus, basal ganglia, and hippocampus) were combined for primary analyses as these aligned with the microdissected deep grey region used for nCounter assessment. Pyknotic nuclei counts displayed a right-skewed distribution and were log transformed prior to analysis. Data were presented as mean with standard deviation (SD). Evidence of synergistic global and regional neuroprotection by Az and Epo was assessed using four theoretical models – combination subthresholding, highest single agent, response additivity, and Bliss independence (*40*). With synergy via combination subthresholding, each drug alone does not produce a significant effect, but the combination does. The “highest single agent” (HSA) model determines the best response obtained using any of the single agents, and this metric is used as a reference point to compare whether the combination provides a larger effect than the HSA. In response additivity, the effect of each drug individually (delta compared to the untreated group) is linearly combined, and synergy suggested if the combination treatment has a larger effect than this combination. Finally, Bliss independence predicts the expected effects of drug combinations based on the assumption that the drugs act independently of each other. The combined effect of two drugs is calculated as if each drug acts on its own without influencing the other’s action. Bliss independence is particularly useful in identifying whether the observed effects of drug combinations are due to true synergistic interactions, though it assumes that both drugs are protective individually.

Comparison of proportions of regional VAMPIRE SMs across groups was performed using a Chi-Square test followed by logistic mixed effects models for each SM individually. When comparing the effect of group on SM distributions across multiple regions, a Mantel-Haenszel (multivariable Chi-square) test was used to adjust for the effect of brain region. nCounter data were examined as log2-fold changes compared to OGD both globally and regionally and compared using a t-test. Volcano plots and heatmaps of significant transcriptomic shifts were generated in Prism (version 10, GraphPad Software, San Diego, USA) and Microsoft Excel, respectively. To reduce the dimensionality of the nCounter data, principal component analysis (PCA) of normalised nCounter expression level was performed. The principal components (PC) that explained 95% of the variance of the data were used to explore the relationship between nCounter transcript groupings and regional and global cell death and microglial SMs. The relationships between microglial morphology and nCounter PCs with microglial SMs and region-specific neuroprotection were assessed using graphical network models. Graphical network analysis involves extracting significant relationships from a precision matrix of inter-related variables, which allows for the identification of important relationships after taking into account how all the other variables are related (*41*). To determine significant relationships in the network we used the method described by Williams and Rast. The precision matrix was constructed using a maximum likelihood estimation (MLE) method and significant relationships were determined using Fisher Z-transformed 95% confidence intervals (*42*). After graphical network analysis, the nCounter transcripts that most contributed to PCs that displayed significant relationships with outcomes of interest were then identified based on their relative contributions and loadings within each PC. To examine potential signatures of emergence, we employed a Bayesian Additive Regression Trees (BART) machine learning model to predict mean log-transformed and normalised gene expression levels in the AZ*Epo group using expression levels in the control, OGD, Epo, and Az groups. 10-fold cross-validation as used to determine pre-validated predications for all 255 nCounter transcripts globally, as well as within each region, using standard BART parameters as previously described.(*43*) Regressing predicted expression levels against measured expression levels in the Az*Epo group, transcripts whose expression was outside the 95 and 99 percent intervals were identified and their difference in predicted versus measured average expression determined as potential candidates for Az*Epo synergy. All analyses were performed in R version 4·4·0 in the RStudio environment. P-values <0·05 were considered statistically significant.

## Results

### Az and Epo result in synergistic global and regional neuroprotection

To investigate the neuroprotective potential of Az and Epo, we used LDH release and percentage of pyknotic nuclei to assess global and regional cell death, respectively. LDH release was increased by OGD and significantly reduced by the Az*Epo combination (p=0·006) but not either treatment individually (**Figure 1A**). Examples of pyknotic nuclei are shown in **Figure 1B**. When analysed across all assessed regions, OGD significantly increased the number of pyknotic cells compared to control (p<0·0001), and this was significantly decreased only by the Az*Epo combination (p=0·01, **Figure 1C)**. In the white matter, the number of pyknotic nuclei decreased with Az (p=0.03, **Figure 1D)**. In the combined deep grey matter regions of the hippocampus, thalamus, and basal ganglia, cell death decreased with Az*Epo (p=0.006, **Figure 1E**), though the response had high variability and was not significantly protective within each region individually except for the hippocampus **(Figure S2A-C).** Az alone was the only treatment that was significantly neuroprotective in the cortex (p=0.004, **Figure 1F**).

**Figure 1.**
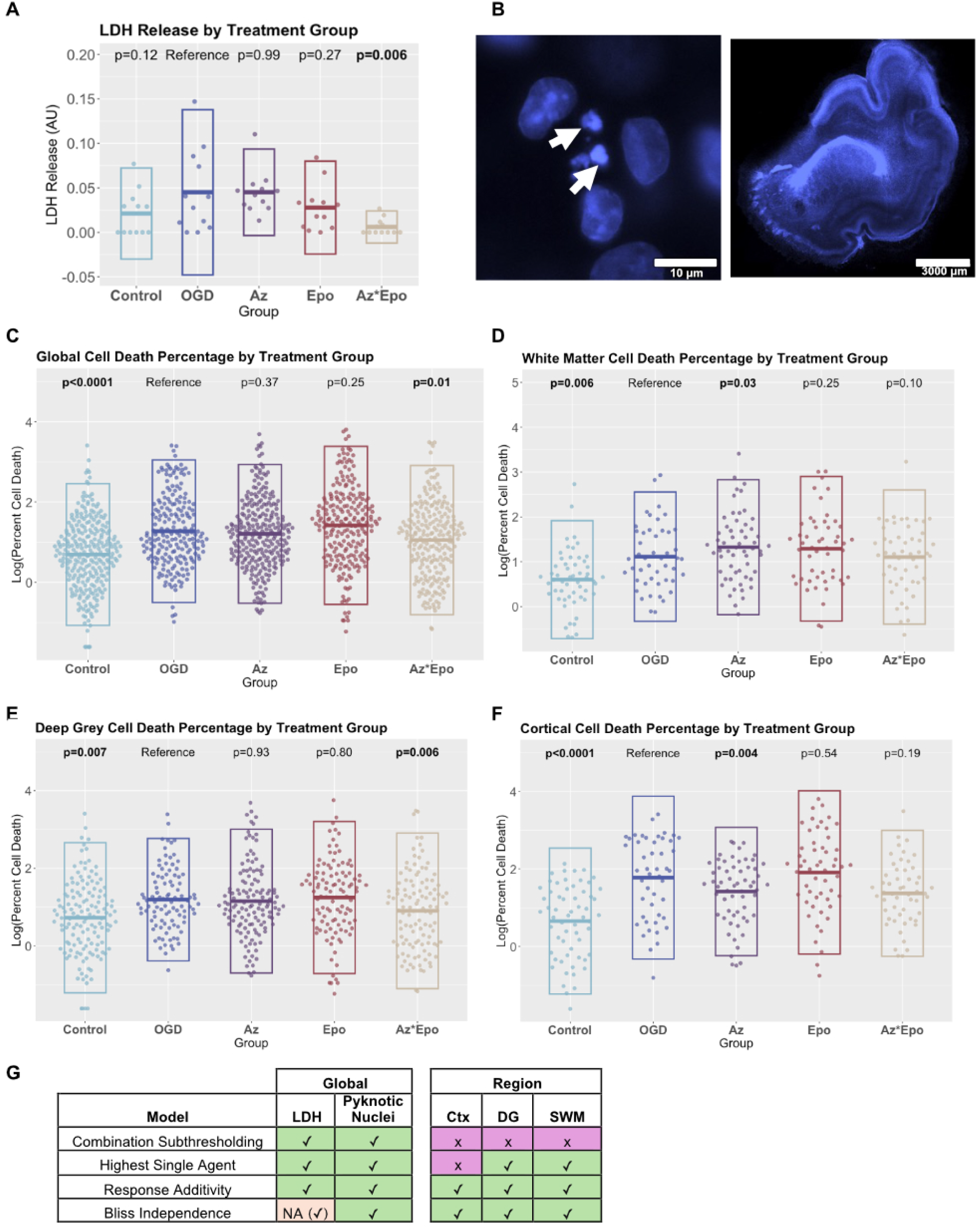
Global cytotoxicity and regional cell viability in response to injury and treatments. (A) Cytotoxicity level quantified by lactate dehydrogenase (LDH) release into the culture media at treatment endpoint (n = 12 slices per group). The combination of Az*Epo decreased LDH levels significantly to the control level, whereas the monotherapy groups saw no effect. (B) Representative immunofluorescence image of pyknotic nuclei (white arrows) indicating cell death (left) and a tile-scan of DAPI-stained whole hemisphere brain slice (right). Log-transformed cell death percentage by treatment groups (C) globally, in the (D) white matter, (E) deep grey matter, and (F) cortex are shown. Each point represents one of n = 3-4 individual images per region per group from n = 9 slices. Boxes show mean with SD. Bolded p-values indicate significant differences compared to the untreated OGD group performed with linear mixed effects models with fixed effects of region and random effect of slice. (G) Synergism of Az*Epo combinatorial treatment evaluated by LDH global cytotoxicity and regional cell death percentage compared to individual therapies. Green ticks indicate evidence for synergy under that model in that region/assessment type. Purple Xs are displayed when no evidence of synergy is seen. NA with a tick indicates where the conditions of Bliss independence were not fully met (Epo did not display any effect on its own) but synergy was suggested according to the model.

Using four well-described statistical models to assess treatment synergy, Az*Epo was synergistically neuroprotective at reducing LDH release and pyknotic nuclei counts according to all models, though one assumption of the Bliss independence model (that both treatments display some degree of protection individually) was not met as either Az or Epo had an average pyknotic nuclei count above that seen in the OGD group **(Figure 1G)**. Across the cortex, deep grey matter, and white matter, evidence of synergistic neuroprotection by Az*Epo was seen according to response additivity and Bliss independence. Within the individual regions of the deep grey matter, some evidence for synergistic neuroprotection was seen in all regions **(Figure S2D)**.

### Az, but not Epo or Az*Epo, reverses OGD-induced loss of microglia

The OGD group showed significantly lower Iba-1+ cell numbers compared to the control group in all regions (**Figure 2A-D**). Treatment with Az significantly increased microglial counts globally (**Figure 2A**, p=0·018) and in the deep grey matter (**Figure 2C**, p=0·012). Representative immunofluorescence images from the median image in each group show nonsignificant increases in Iba-1+ cell counts in the white matter and basal ganglia in the Az group compared to OGD (**Figure 2E**). The cell numbers in these regions became comparable to the control group. However, neither Epo nor the Az*Epo combination significantly changed microglial numbers relative to OGD. No significant changes in Olig2+ cells were seen by region or treatment group either globally (**Figure S3A**) or within the subcortical white matter (**Figure S3B**).

**Figure 2.**
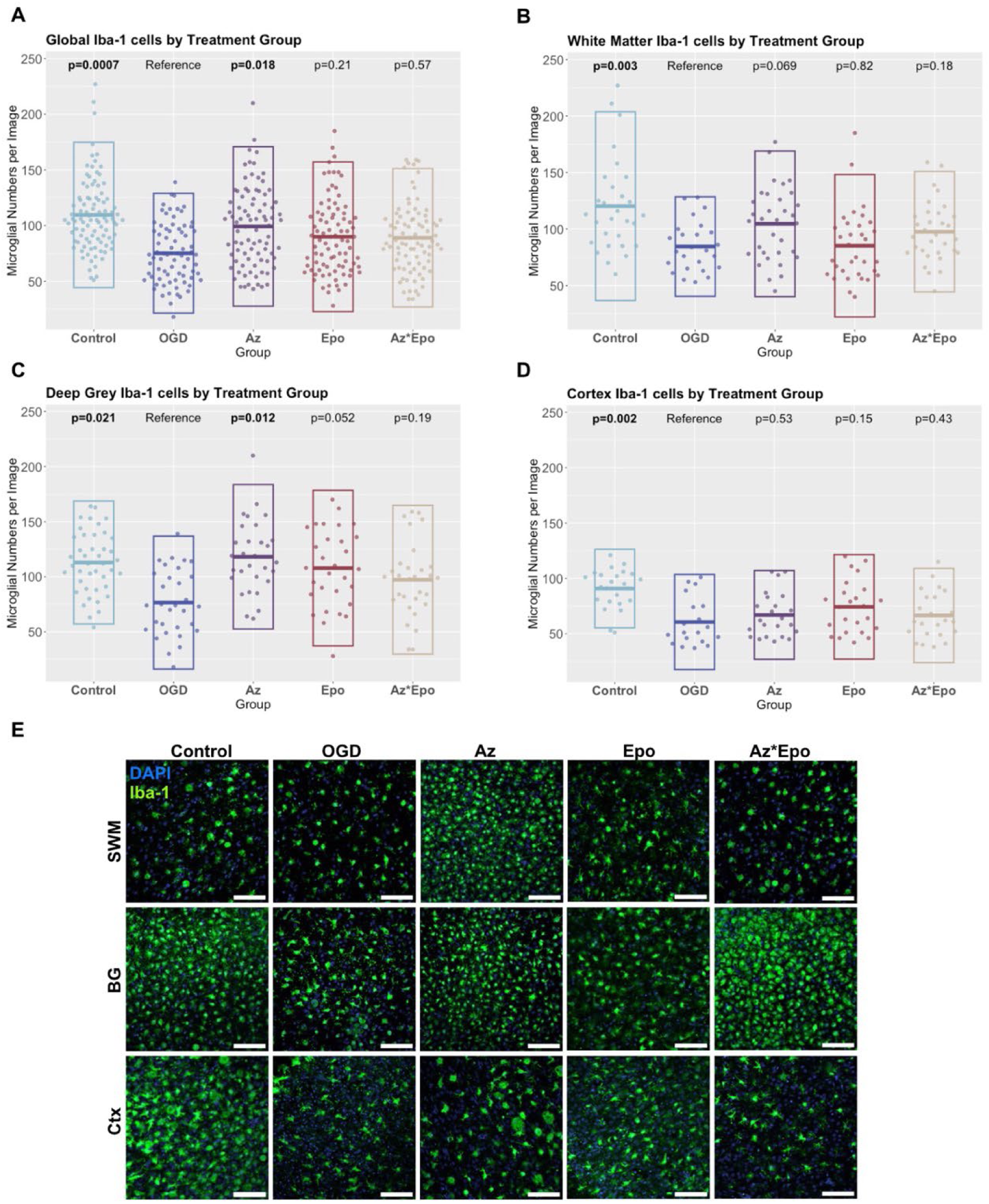
Regional microglia cell number response to OGD and treatments. Iba-1+ cell counts by treatment groups (A) globally, in the (B) white matter, (C) deep grey matter, and (D) cortex are shown. Each point represents one of n = 3-4 individual images per region per group from n = 9 slices. Boxes show mean with SD. Bolded p-values indicate significant differences compared to the untreated OGD group performed with linear mixed effects models with fixed effects of region and random effect of slice. (E) Representative immunofluorescence images stained with Iba-1 for microglia by treatment groups in the subcortical white matter (SWM), basal ganglia (BG) in the deep grey matter, and cortex (Ctx). Scale bars are 100μm in images.

### Neuroprotection involves augmentation of certain microglial responses to injury

*In vitro* and *in vivo* experimental data have indicated early and persisting phenotypic changes in microglia after neonatal HI (*44, 45*). In addition to changes in cell number, morphological shifts in microglia can provide insights into microglial phenotypic responses to injury and treatment. As we measured several interrelated microglial morphology parameters, we first used graphical network analysis to determine significant relationships with OGD and treatment after accounting for relationships between the different parameters (**Figure 3**). Comparing OGD to control, OGD appeared to significantly increase cell circularity and eccentricity, where larger eccentricity values imply a more elliptical shape. OGD also decreased solidity and major axis length, where smaller solidity values and major axis lengths suggest a more ramified and condensed morphology (**Figure 3A, Table S1**). Comparing the treatment groups to OGD, all three treatments appeared to reverse the decrease in solidity and major axis length seen with OGD (**Figure 3B-D**), and Az*Epo reversed the increase in eccentricity (**Figure 3D**). Az alone further increased cell circularity compared to OGD (**Figure 3C**). Similarly, when comparing microglial morphology in regions that experienced neuroprotection by any of the treatment groups to those that did not, neuroprotection was associated with increased circularity and decreased cell area (**Figure 3E**). This suggests that neuroprotection may occur partly by augmenting certain initial microglial morphological responses to injury.

**Figure 3.**
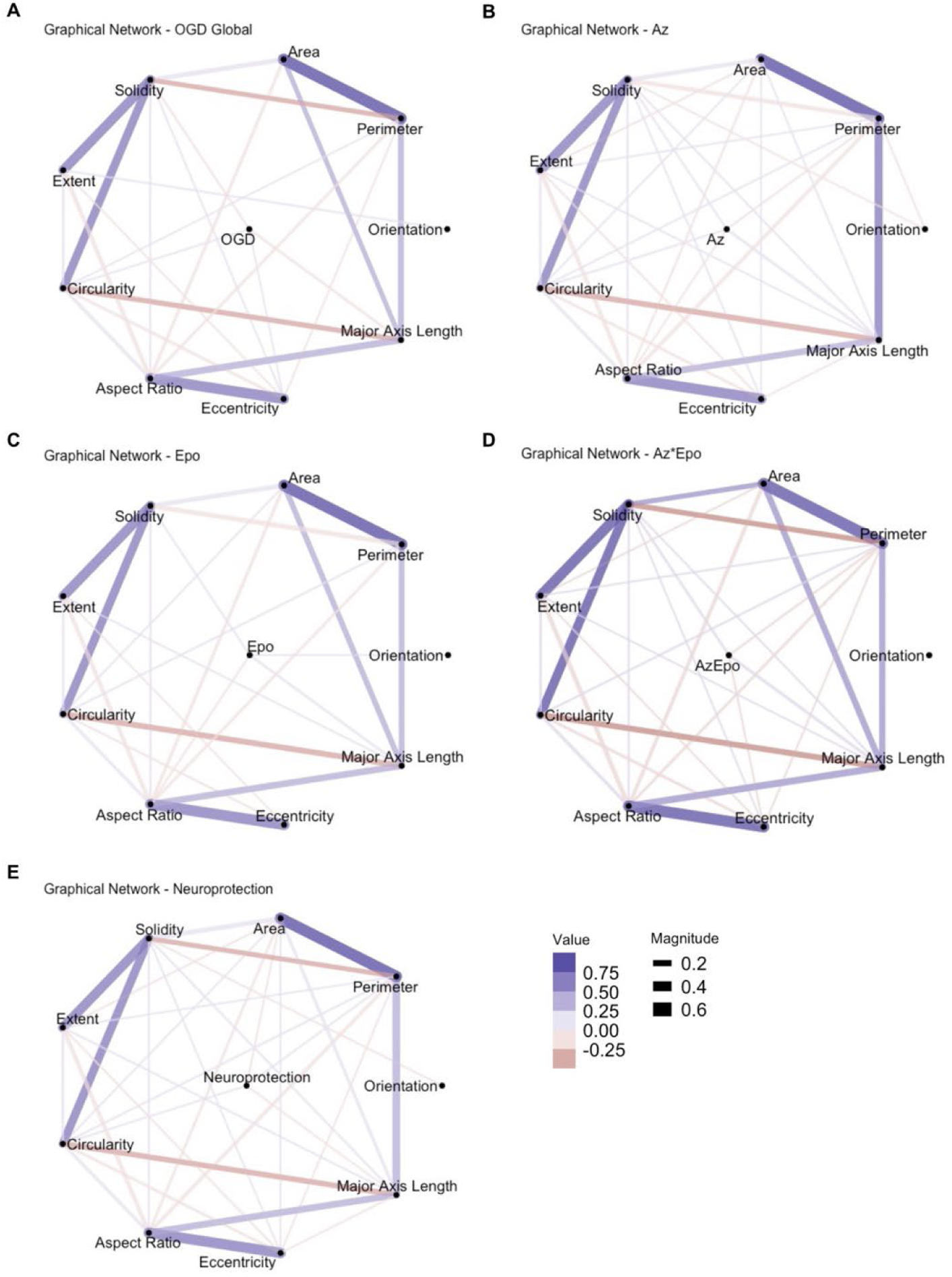
Microglial morphological parameter trends in relation to OGD, treatments, and neuroprotection. Graphical network analyses of microglial morphological parameter changes in response to (A) OGD, (B) Az, (C) Epo, and (D) Az*Epo treatment conditions globally are shown. The morphological parameters including area, perimeter, orientation, major axis length, eccentricity, aspect ratio, circularity, extent, and solidity were quantified from microglial cells captured from n = 3-4 individual images per region from n = 9 slices in each group. (E) Microglial morphology parameter trends in regions experiencing significant neuroprotection (by any treatment) are shown. Lines between two points indicated a statistically significant correlation after accounting for the other relationships in the network. Purple lines indicate significant positive trends between the morphological parameters and conditioning. Red lines indicate significant negative trends between the morphological parameters and condition. The thickness of the lines displays the strength/magnitude of the relationship.

### Microglial shape mode shifts identify regional neuroprotection by Az and Az*Epo

To characterise microglial morphological states and their distributions by region and treatment, we clustered individual microglia into distinct populations through PCA **(Figure 4A)** and classification into five morphological SMs using the VAMPIRE pipeline (*17*). We found that regional variability in SM distribution was apparent by treatment group. Globally, Az, Epo, and Az*Epo had significantly lower distribution of SM1 microglia compared to OGD (**Figure 4B**). The combinatorial Az*Epo treatment group in addition showed a significantly higher proportion (31%) of SM5 microglia compared to the OGD group (22%). We further explored whether these proportions differed by region where treatments were and were not neuroprotective. While a significant decrease in SM1 remained consistent across all treatment groups regardless of neuroprotection, the increase in SM5 with treatment became more apparent in both the Az and Az*Epo group only when neuroprotection was not seen (**Figure 4C**), which was primarily in the thalamus.

**Figure 4.**
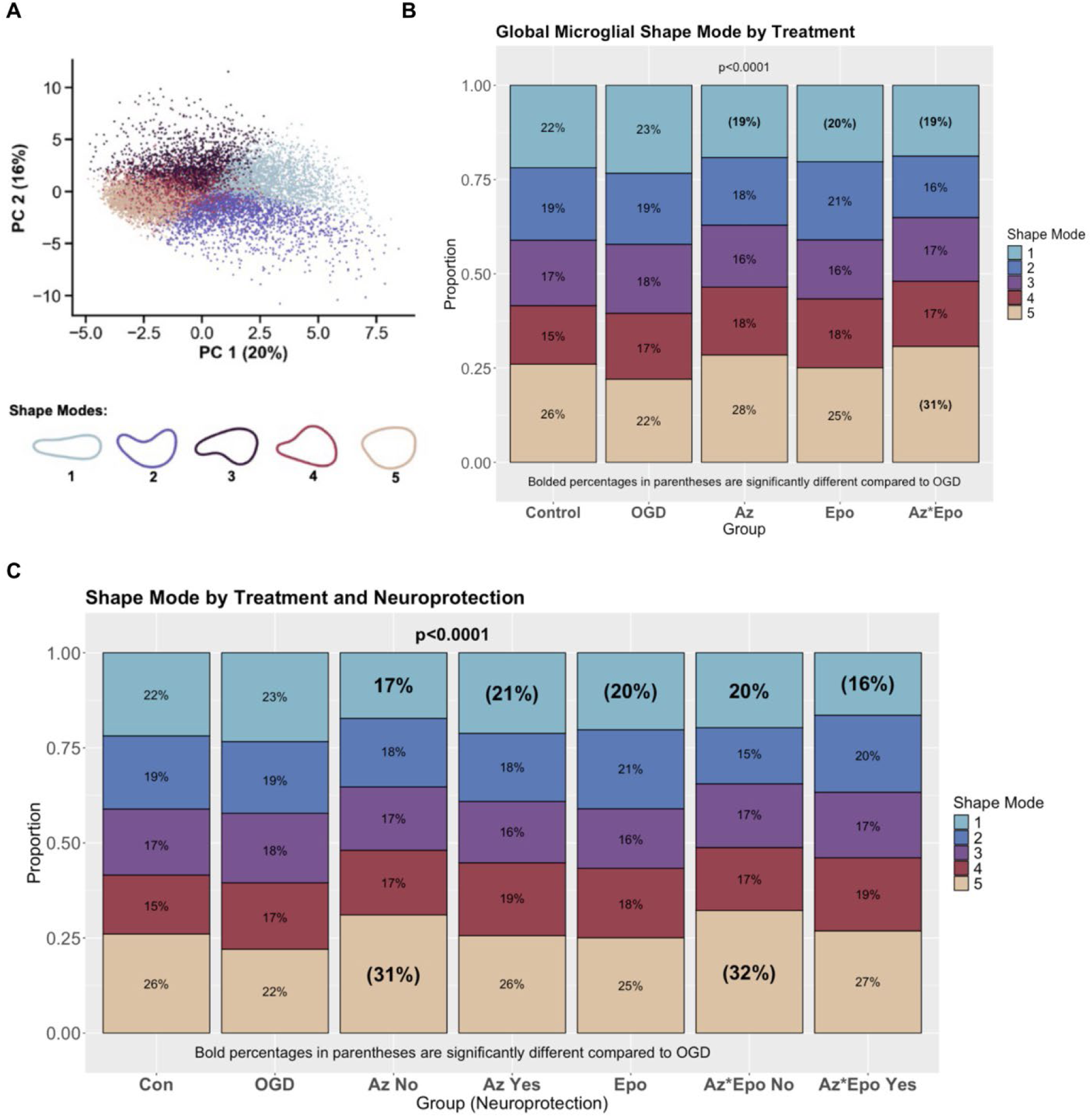
Regional microglia morphology changes by shape mode in response to OGD and treatments. (A) PCA plot shows separation of five microglial morphological groupings using the first two derived PCs. (B) Proportions of five different microglial SMs by treatment groups are shown globally. (C) Proportions of different SMs by treatments in regions that were and were not neuroprotective (based on pyknotic cell counts). Bolded percentages in parentheses indicate SM proportion that is significantly different compared to the untreated OGD group. Data was gathered from n = 3-4 individual images per region per group from n = 9 slices.

We also used graphical network analyses to explore how SMs related to morphological parameters discussed above. SM1 microglial phenotype was positively correlated with aspect ratio, where a higher value suggests the presence of a more elongated cell body **(Figure S4A).** SM1 was also positively related to solidity, where greater solidity is associated with a less ramified cell body. The primary notable relationships between microglia parameters and both SM2 **(Figure S4B)** and SM3 **(Figure S4C)** were a positive association with eccentricity and a negative association with aspect ratio. No notable primary drivers of SM4 were identified **(Figure S4D)**, which received a small but significant contribution from most of the parameters examined. We found that the SM5 phenotype was negatively correlated with area and eccentricity, where the eccentricity value of close to 0 presents the shape of a circle and close to 1 presents an elongated ellipse **(Figure S4E).** Combining all the microglial shape mode and parameter trends, the distribution of elongated, ramified, and rod-shaped microglial SM1 morphology was decreased by all treatments regardless of neuroprotection. Only in regions where Az and Az*Epo did not have neuroprotective effects, rounded-shaped microglia with less cell area coverage (SM5) were increasingly observed compared to OGD.

### Lack of regional neuroprotection is associated with persistence of reactive pro-inflammatory microglia

To reduce the dimensionality of the nCounter data, we performed PCA on the normalised transcript levels and selected the seven PCs that explained at least 95% of the variance of the nCounter data (**Figure S5**). We then aligned nCounter PCs by region and treatment to the corresponding slices with Iba-1 imaging. As SM1 and SM5 appeared to be particularly responsive to OGD and treatment, we focused on nCounter PCs associated with the relative proportion of those SMs by region and treatment group. Using graphical network analyses, PCs 2 and 5 appeared to be most strongly associated with SMs 1 and 5 in a reciprocal manner (**Figure 5A and B**). PC2 was negatively associated with SM1 and positively associated with SM5, and PC5 was positively associated with SM1 and negatively associated with SM5. We then explored the top 25 genes contributing to PCs 2 and 5 based on their contributions and loadings displayed (**Figure 5C)**. Within these genes, some contribution to PC2 appeared to be due to microglia-related genes such as TREM2 and AXL, involved in microglial homeostasis and pathogen defence, respectively. By comparison, PC5 was negatively associated with several markers of microglial reaction and inflammation, including CXCL10, STAT1, and CD86.

**Figure 5.**
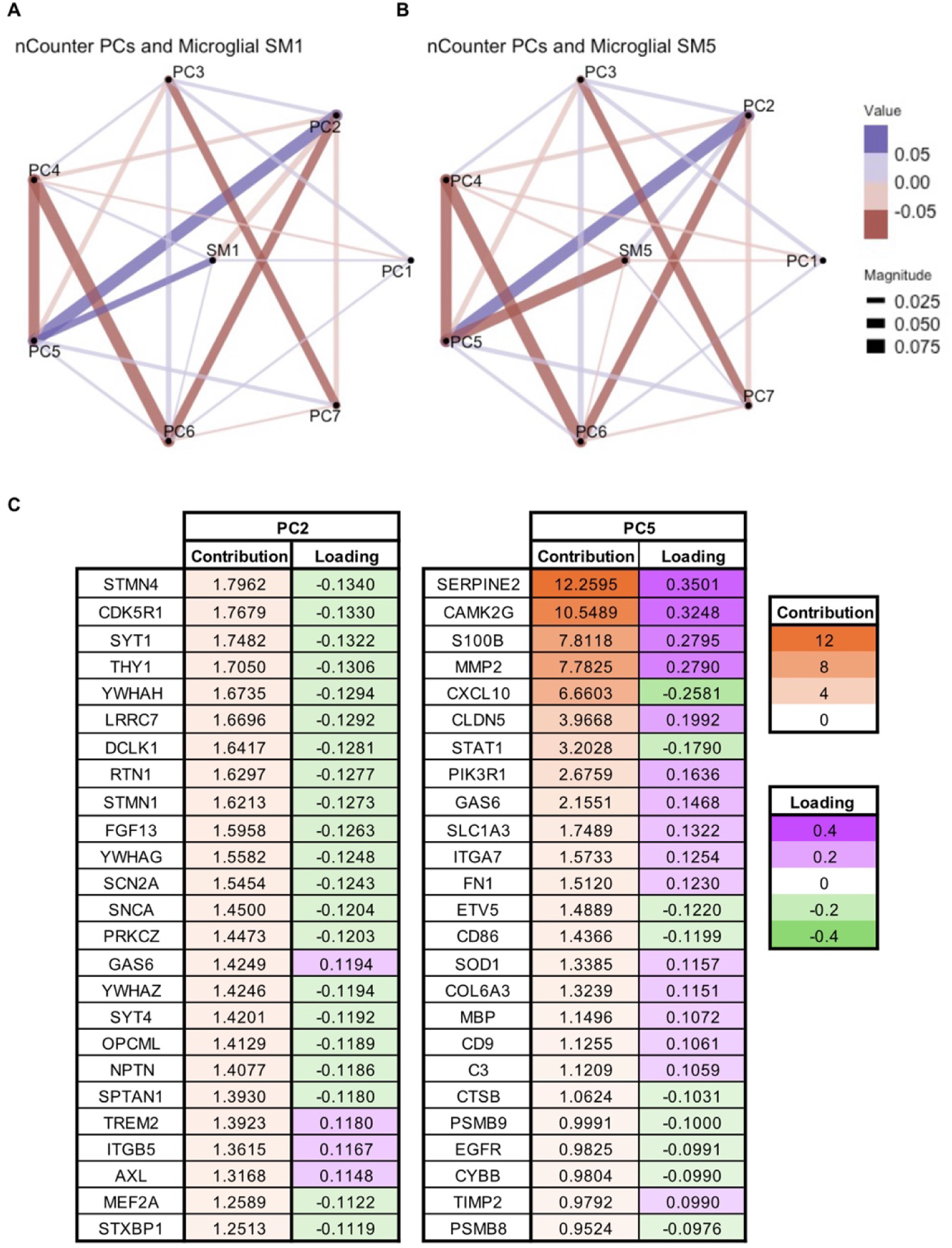
nCounter transcript PCs in relation to microglial shape modes of significant responses to OGD and treatments. Graphical network analyses of seven PCs of nCounter data in relation to microglial (A) SM1 and (B) SM5, which appeared to have significant distribution changes in response to treatments compared to OGD, are shown. In the graphical networks, lines between two points indicated a statistically significant correlation after accounting for the other relationships in the network. Purple lines indicate significant positive trends between the morphological parameters and conditioning. Red lines indicate significant negative trends between the morphological parameters and condition. The thickness of the lines displays the strength/magnitude of the relationship. The thickness of lines in graphical networks imply how positively or negatively correlated the relationships were. (C) Contributions and loadings of top twenty-five contributing genes in PC 2 and 5, from which notable associations with SMs 1 and 5 were observed.

As higher SM5 in regions such as the thalamus not experiencing Az- and Az*Epo-related neuroprotection (**Figure 4C**) would be associated with reduced expression of genes in PC5, and PC5 is negatively associated with pro-inflammatory microglial transcripts, this suggests that microglia in regions not experiencing neuroprotection may have expressed an inflammatory phenotype that was not amenable to modulation by Az or Epo. Relationships between nCounter PCs and microglial SMs 2, 3 and 4, as well as selected microglial morphology parameters are shown in **Figure S6A-C.** PC5 was strongly negatively associated with circularity (**Figures S6D**) and positively associated with perimeter (**Figure S6E**). Full contributions and loadings for all nCounter PCs are shown in **Table S2**.

### Az*Epo synergism results in both augmentation and reversal of specific transcriptomic responses to injury

We then examined relationships between nCounter PCs and responses to OGD and treatment. Comparing OGD to control, OGD was positively associated with PCs 1-6, and negatively associated with PC 7 (**Figure 6A**). When combining the treatment groups and comparing regions that saw significant neuroprotection compared to those that did not, neuroprotection was positively associated with PCs 1, 4, and 5, and negatively associated with PCs 2, 3, 6, and 7 (**Figure 6B**). These relationships were even stronger when focusing on regions that did and did not experience neuroprotection by Az*Epo (**Figure 6C**). In this scenario, regional neuroprotection was positively associated with PCs 1, 2, and 5, and negatively associated with PCs 3, 4, 6, and 7. This pattern of potentially synergistic neuroprotection by Az*Epo was therefore characterised by both augmentation and reversal of specific transcriptomic signatures associated with injury (**Figure 6D**). Treatment with Az*Epo appeared to augment transcriptomic signatures associated with PCs 1, 2, 5, and 7 via further positive or negative associations in the same direction as OGD compared to control, and reversed associations – associations were in the opposite direction as OGD compared to control – signatures associated with PCs 3, 4, and 6.

**Figure 6.**
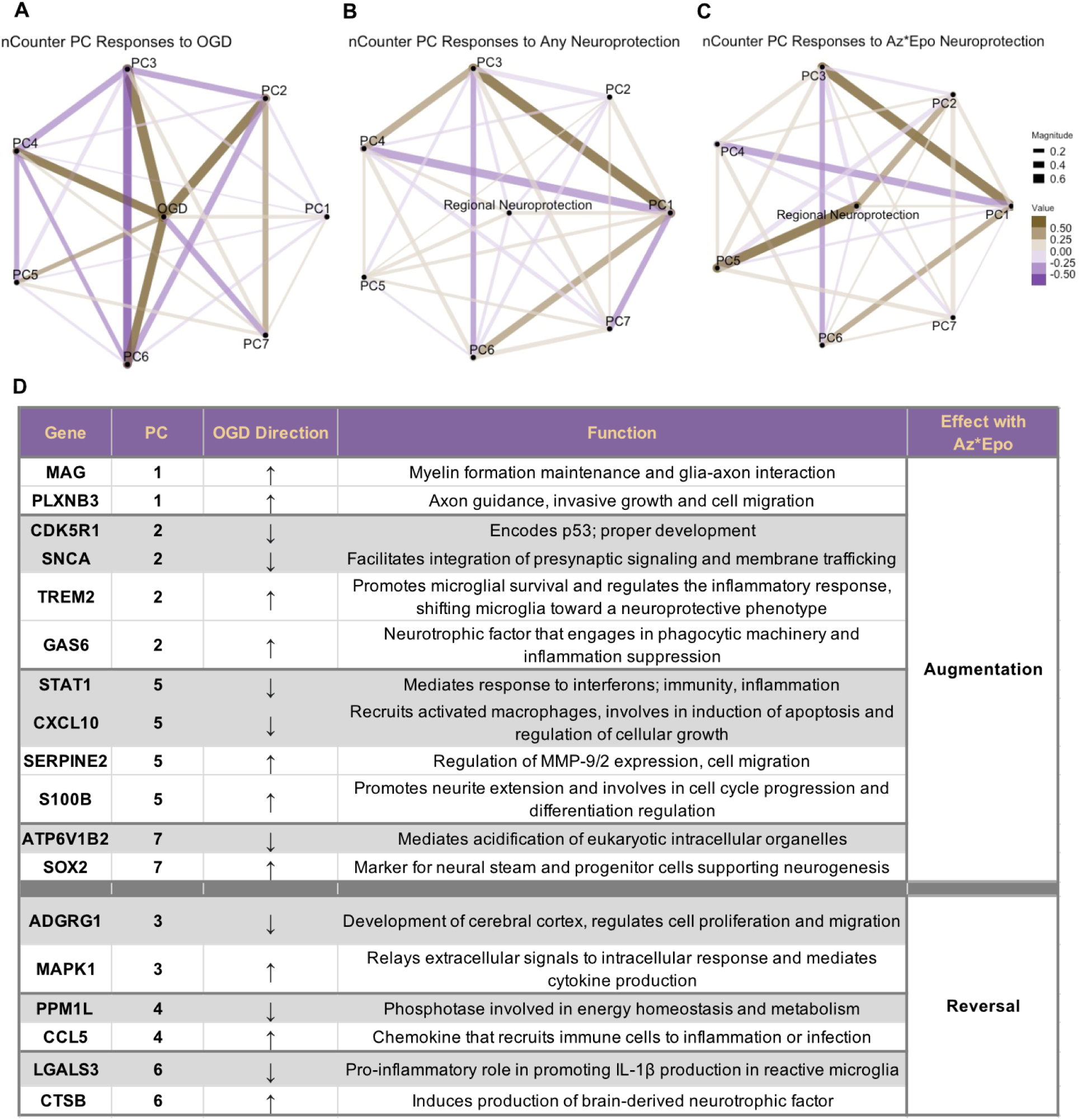
nCounter PCs in relation to neuroprotection mechanisms. Graphical network analyses of seven PCs of nCounter data in relation to (A) OGD injury, (B) any treatment in neuroprotective regions, and (C) Az*Epo combination treatment neuroprotection are shown. Golden lines indicate significant positive correlation between the nCounter PCs and conditions. Purple lines indicate significant negative correlation between nCounter PCs and conditions. The thickness of lines in graphical networks imply how positively or negatively correlated the relationships were. (D) Representative genes from each PC that were interpreted to be augmented or reversed by Az*Epo from previous analysis are listed with whether OGD up- or down-regulated their expression levels and the potential protective mechanisms.

To further investigate the transcriptional changes related to OGD and treatment protection responses, we performed GO enrichment analysis for each nCounter PC to identify gene pathways and network by functionality **(Figure 7, Table S2)**. Combined with the previous results, inferences can be made on the transcriptomic mechanisms that were augmented or reversed by Az*Epo treatment to achieve neuroprotection. The top contributing genes in PC2, including signal transduction genes, synaptic function genes SNCA and SCN2A, and brain development gene CDK5R1, were differentially downregulated in response to OGD (p<0.05, log2 fold change <= -1; **Figure 6D, Table S3 and S4).** Az*Epo neuroprotection seemed to augment these differentially expressed genes (DEGs), which were significantly enriched in regulation of neurogenesis, cell differentiation, and nervous system development **(Figure 7)**. Similarly, top contributing genes in PC5, inflammation and metabolic reactive genes STAT1, SOD2, and CXCL10 also had decreased expression levels in response to OGD with augmentation by Az*Epo treatment. The upregulated expression of genes such as TREM2, GAS6, SOX2, S100B, and SERPINE2 involved in microglial homoeostasis, suppression of inflammation, neurogenesis, neurite extension, and cell migration, respectively, were also further augmented (**Figure 6D**).

**Figure 7.**
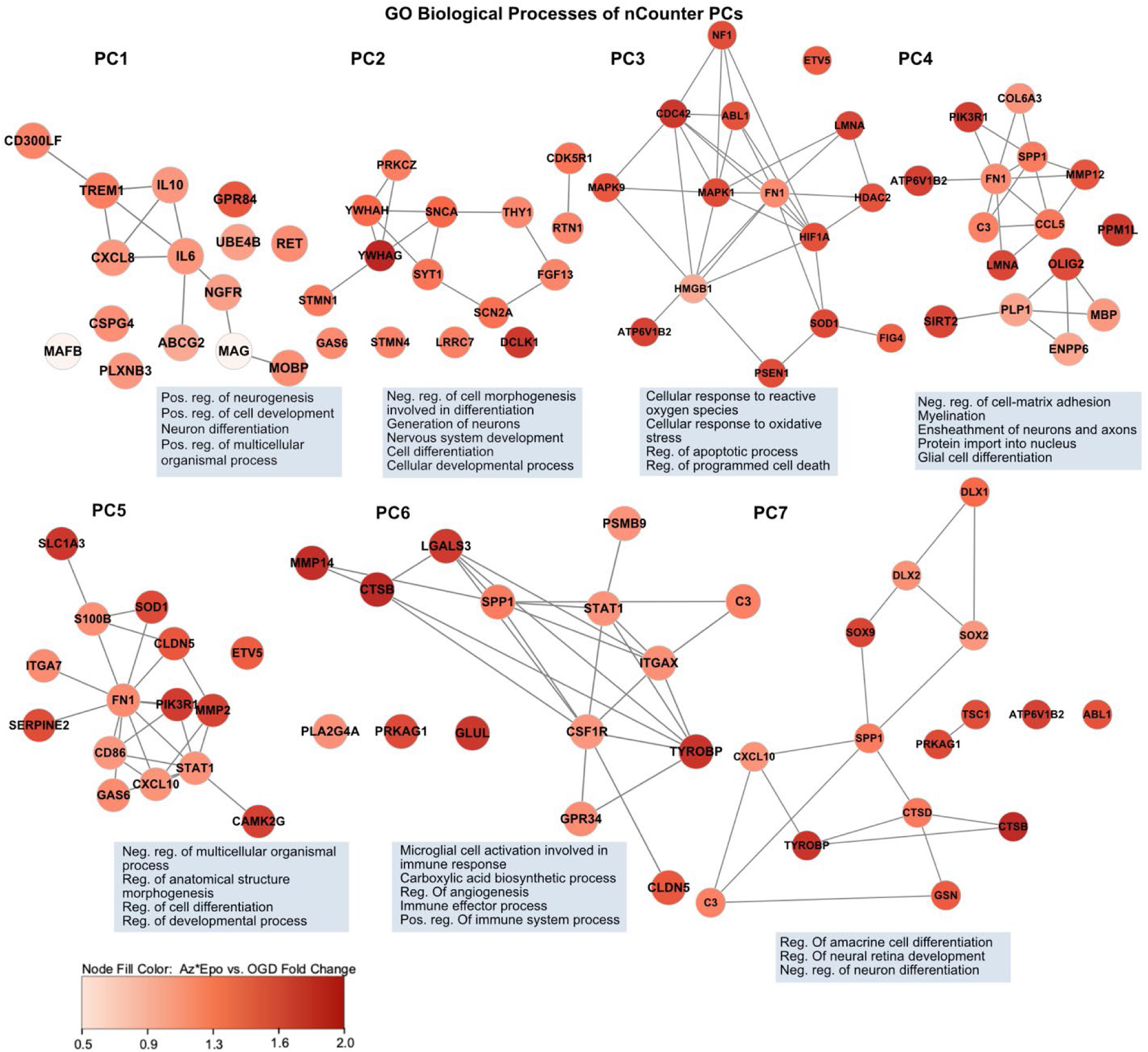
Transcriptional states and expression levels compared to OGD by nCounter PCs. The top fifteen contributing genes in each nCounter PCs are represented in the co-expression networks. The edge represents the co-expression in other transcriptional states. The colour of nodes represents the degree of gene expression level fold change with Az*Epo treatment application post-OGD. The top five most significantly enriched GO terms (cutoff p-value < 0.05) that are associated with signature genes of each PCs are indicated. For GO terms that were semantically similar (e.g., “cellular response to oxidative stress” and “response to oxidative stress”), the less significant GO term of the two was removed.

PC3 genes enriched in regulation of apoptotic and cell death processes, including LMNA responsible for cellular structural integrity, cytoskeletal regulator CDC42, and cell differentiation regulator MAPK1, were downregulated with OGD. These transcriptomic signatures associated with OGD were reversed by Az*Epo **(Figure 6D and 7)**. We also found that the top contributing transcripts to PC6, which was positively associated with OGD and reversed by Az*Epo, were significantly enriched in angiogenesis and immune response-related genes including CTSB, MMP14, and TYROBP. Combinatorial Az*Epo treatment appeared to also reverse OGD-induced changes in expression of PC4, which included enrichment of genes related to several signalling phosphatases and kinases as well as the pro-inflammatory microglia-associated cytokine CCL5.

### Az*Epo suppresses glial activation and inflammation

We then explored the DEGs in brain regions for each treatment condition. Globally, 92 genes showed differential expression post-OGD while regionally OGD resulted in 145 DEGs in the white matter, 77 DEGs in the deep grey matter, and 64 DEGs in the cortex **(Figure 8A, Table S3).** In the white matter, where the greatest number of DEGs were identified in response to OGD, OGD was associated with upregulation of glial differentiation and downregulation of several pathways associated with neuronal regulation, differentiation, and survival **(Figure 8C).** Globally, only four genes related to macrophage differentiation (MAFB), myelin sheath formation and maintenance during nerve regeneration (MAG), and cell cycle regulation (AHCYL1 and CDKN1A) were differentially regulated across all three treatment conditions (Az, Epo and Az*Epo). MAFB is also linked to the GO term “rhombomere 5 development”, which involves neuro-epithelium patterning during early hindbrain development (*46*) **(Figure 8B)**. We investigated phenotype profiles specific to treatments in white matter where most genes were differentially influenced post-OGD. Genes related to blood-brain barrier permeability (CLDN5), cell-cell adhesion and interaction (ITGA7, PSEN1), DNA repair facilitation (PARP1), and metabolic activity (SOD2, PPM1L) were differentially regulated by Az **(Figure 8B, Table S3 and S4).** Genes related to synaptic homeostasis and plasticity (COL6A3, SERPINI1), peripheral nerve regeneration (NTF3), and neural morphogenesis and connectivity (ARHGEF10) were regulated by Epo treatment. In the deep grey matter, C5AR1, an inflammation-responsive gene, was differentially regulated by Az treatment alone, in addition to CDKN1A and CD40. Epo treatment was associated with regulation of BCL2L11, responsible for cell homeostasis and apoptosis regulation, and EGFR.

**Figure 8.**
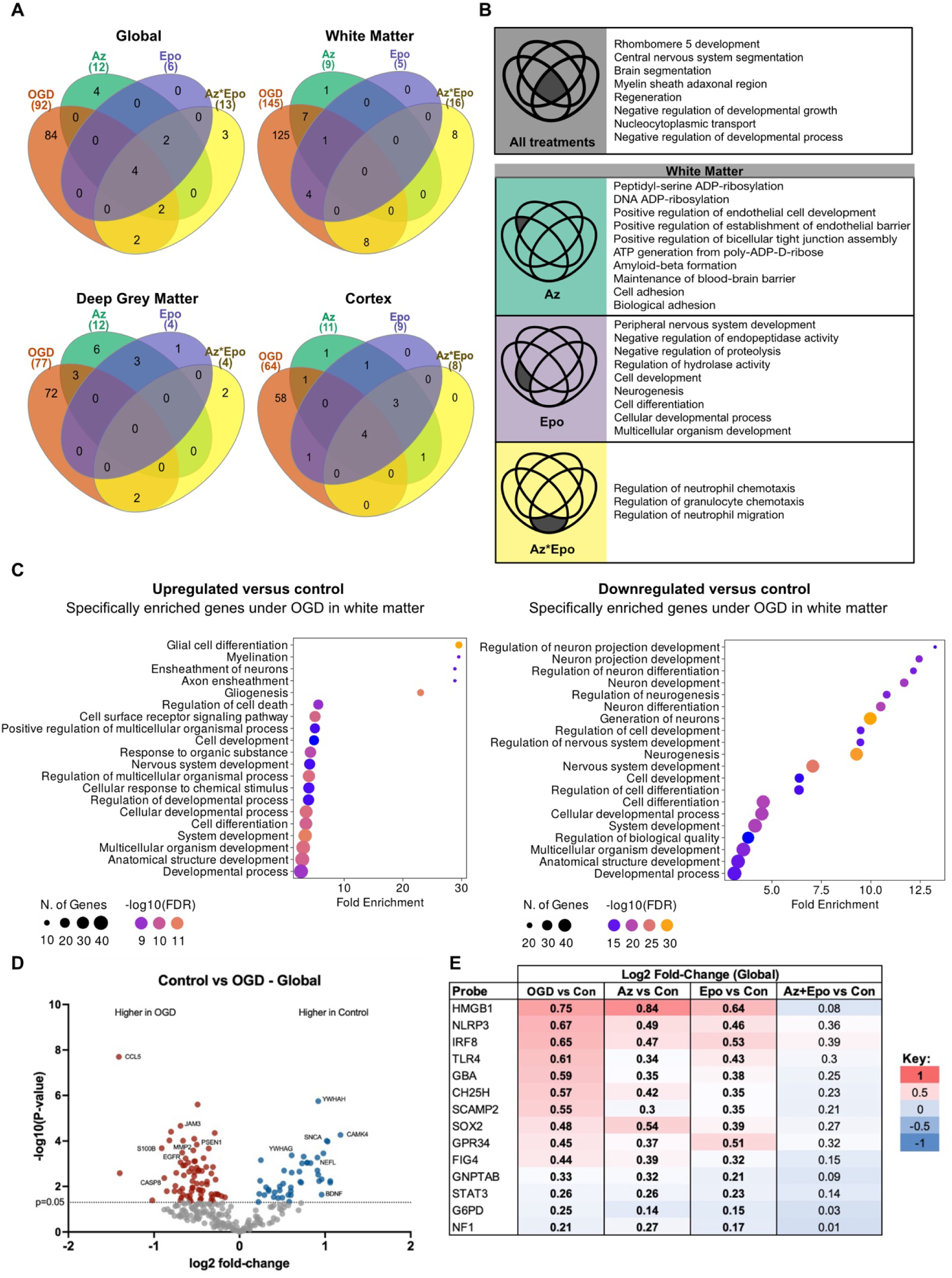
Regional transcriptional state changes to OGD and treatments and emergent transcriptomic signatures of global synergistic protection by Az*Epo. (A) Signature genes represented in the Venn diagrams were determined by comparing gene expression levels of OGD versus control, and all treatments versus OGD globally and in the white matter, deep grey matter, and cortex with p-values <0·05. (B) The top ten significant GO terms that are associated with signature genes identified in (A) are listed. Dark grey regions in the Venn diagrams indicate gene set that was regulated by the specific treatment depicted. Significant GO terms were ranked by fold enrichment. (C) The representative enriched GO biological process of up- and down-regulated genes under OGD compared to control are shown. -log10 (FDR) and gene counts are indicated in the dot plots. (D) Volcano plots of globally differentially expressed genes after OGD. Coloured dots represent genes that were significantly differentially expressed comparing with the control group with p value < 0·05. Positive log_2_ fold-change (red) indicates upregulated expression after OGD, and negative log_2_ fold-change (blue) indicates downregulated expression. (E) The heatmap shows log_2_ fold-change globally for each group compared to control. Bolded values are biomarkers that are significantly differentially expressed compared to control (p<0·05, t-test). The genes that were normalised by Az*Epo but not by Az or Epo individually are shown. N = 6 slices per region from each treatment group.

An inflammatory signature was prominent with genes regulated by the combinatorial Az*Epo treatment. This was shown by the top GO terms, including regulation of neutrophil migration and chemotaxis, which are activated as part of the innate immune response. Genes of macrophagic and proteolytic activities (MARCO, CTSD, PSEN2), neuronal differentiation (TIMP2), and other glial cell activation and regulation (EGFR, TSPO, JAM3) were particularly differentially regulated by Az*Epo. Overall, regional transcriptional trends in both the white matter and deep grey matter suggested that the Az*Epo treatment profile was associated with suppression of inflammation and glial cell activation **(Figure S7)**.

### Transcriptomic responses to Az*Epo suggest emergent synergism

While neuroprotection was not uniformly associated with reversal of transcriptomic responses to OGD, we finally explored whether there were any signatures uniquely associated with Az*Epo when applied in combination. As Az*Epo most strongly displayed synergy by reducing LDH release, we compared first global nCounter transcripts in OGD versus control (**Figure 8D**), and then extracted transcripts that were significantly differentially regulated by OGD and normalised by Az*Epo but not either drug individually (**Figure 8E**). Fourteen transcripts satisfied these criteria, with the top two being HMGB-1 and NLRP3. HMGB-1 is a critical regulator of transcription and acts as a damage-associated molecular pattern when it is released during cell death. NLRP3 is an inflammasome critical to pyroptotic cell death and widely implicated in a range of neonatal brain injuries. The next two most synergistically normalised transcripts were IRF8 and TLR4, both of which are upregulated in reactive and pro-inflammatory microglia. Therefore, several putative pathways exist by which emergent synergistic properties of neuroprotection by Az*Epo may occur, and these may not necessarily be predicted by responses to each drug individually. Transcriptomic signatures associated uniquely with Az*Epo treatment in the white matter included the reversal of upregulated macrophage response receptor MARCO, and an increase in the downregulated C3, which is involved in microglial activation and neurogenesis (**Figure S8A-C**). In the grey matter, Az*Epo synergistically downregulated IL10 and PLXNC1, suggesting a reversal of inflammatory response to injury, and normalised a suppression of IGF1, which is critical in promoting neurogenesis and myelination in the preterm brain (**Figure S8D-F**).

As an alternative approach to determining whether emergent signatures were evident from the transcriptomics data, we used the non-linear machine learning algorithm BART to predict gene expression level in the Az*Epo group using expression levels from the control, OGD, Epo, and Az groups (**Figure 9A-D**). Genes whose measured transcript expression levels were outside the 95 and 99 % prediction intervals for that region were considered candidates for signatures of emergence, and their absolute difference in normalised expression (Z-score of log-transformed expression) were determined (**Figure 9E**). Notable genes overexpressed in Az*Epo above the 99 % prediction interval were TIMP2 globally as well as in the cortex and deep grey matter, and ADGRG1 in the cortex and white matter. TIMP2 is involved in plasticity and regulation of the extracellular matrix, and ADGRG1 is a G-protein coupled receptor involved in brain cortical patterning. Genes underexpressed in response to Az*Epo compared to predictions included PARP1 globally, MAPK14 in the cortex, and FABP5 in the deep grey. PARP1 is a critical regulator of DNA repair but its overactivation can trigger the NAD+-dependent cell death *parthanatos*. MAPK14 is a regulator of stress responses, and FABP5 is implicated in several inflammatory disease processes. Together, these results suggest that synergistic neuroprotection by Az*Epo involves emergent upregulation of plasticity processes while downregulating stress, inflammatory, and cell death-related responses.

**Figure 9.**
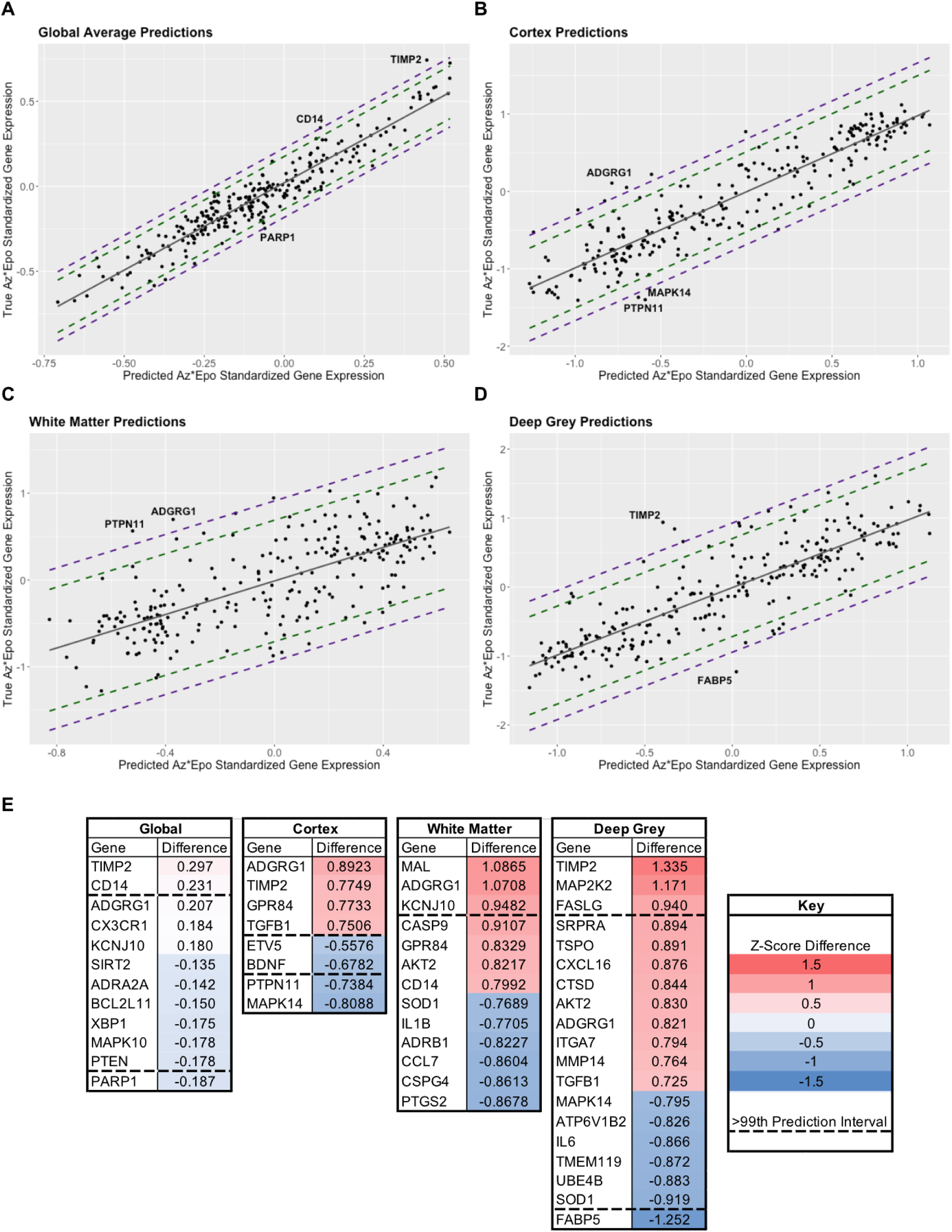
Signatures of emergence from machine learning predictions of Az*Epo nCounter expression. Global (A), cortical (B), white matter (C), and deep grey (D) plots of pre-validated predictions of normalised gene expression relative to measured expression level of all 255 nCounter transcripts. Solid grey line is line of equivalence, with 95 % and 99 % prediction internals in dashed green and purple lines, respectively, generated from predictions using the BART machine learning algorithm. (E) Heatmaps of absolute difference between predicted and measured mean log-transformed and normalised gene expression for transcripts outside of the 95 % prediction intervals globally and for each region. Numbers indicate Z-score difference (number of standard deviations of absolute difference in expression level) between the predicted and measured expression levels. Transcripts above and below the dashed lines are those outside the 99 % prediction level and therefore the strongest candidates for signatures of emergence.

## Discussion

In this study, we developed an extremely preterm-equivalent ferret OWH brain slice OGD injury model to investigate injury responses and the neuroprotective effects of Az, Epo, and their combination, focusing on global and regional cellular analyses and targeted transcriptomics. We first demonstrated overall cellular changes to Az, Epo, and Az*Epo in regions where injury is commonly seen in preterm infants. Combination treatment significantly reduced global cytotoxicity after OGD, which was not seen with monotherapy. This is in line with the recent PENUT trial where Epo alone was not neuroprotective (*25*), but underscores the potential of Epo as part of a therapeutic cocktail. Az alone and Az*Epo decreased pyknotic cell death in multiple regions, with evidence of synergistic neuroprotection by Az*Epo according to multiple models (*47, 48*). Demonstration of synergism in preclinical studies has driven clinical evaluation of drug combinations in cancer and infectious diseases (*40, 49*), but this has not yet been done routinely in the neurosciences. By quantifying synergism using multiple models built upon mathematical frameworks describing different definitions of additivity, we gain more accurate and reliable insights into drug interactions. While synergy can therefore be considered a theoretical construct, several aspects of the cellular and transcriptomic responses to Az*Epo suggest robust and sometimes emergent (e.g., not predictable based on responses to each treatment individually) synergistic effects associated with neuroprotection.

Microglia are the resident phagocytic cells in the central nervous system and their morphologies and functions are closely linked to reflect microenvironmental changes. Functionally, microglia engage in surveillance and homeostasis, including phagocytosis during axonal and synaptic pruning (*50*), promotion of neurogenesis through cytokine secretion (*51*), and neuronal support through the release of neurotrophic factors. In this study, we captured morphological and phenotypic changes in microglia to probe cellular responses to injury and treatment. Overall, OGD decreased microglial counts compared to control across all regions, consistent with our previous findings (*17*). We also found that Az treatment increased microglial counts globally and in the deep grey matter, but this did not occur when Az was combined with Epo. In the deep grey matter, transcriptomic data showed that EGFR, an important growth factor receptor involved in the biochemical regulation of microglia migration and activation (*52*), was significantly upregulated with Az treatment but not the other treatment groups, which may partly explain this finding. However, microglial expansion did not appear to be necessary for much of the neuroprotective effect of Az when applied in combination with Epo.

In our study, OGD increased circularity and decreased solidity and major axis length, from which a more circularly condensed and hyper-ramified or microglial state can be suspected. This finding is partially consistent with what we have reported in the term-equivalent ferret model that showed activated or ameboid microglial morphology post injury (*17*), with decreased area coverage and perimeter and increased cell circularity (*53*). All treatments regardless of neuroprotection decreased the distribution of SM1 morphology that most resembled rod-shaped microglia, a less commonly reported population distinct from the classic ramified, activated and ameboid morphological states (*53*). A rod-like morphology has previously been suggested to occur as a transient state during the de-ramification process (*54*). Microglia in rod shapes may also form in conjunction with neuronal pathology during the initial phase of activation (*55*). The rod microglial morphology appears to be at least partly controlled by the CX3CR1/CX3CL1 signalling pathway (*55*). In our study, Az lowered CX3CR1 expression after OGD in the white matter, but Epo did not. Importantly, the Az*Epo combination treatment group consistently brought down CX3CR1 levels to control levels across all regions. This suggests that Az*Epo may be neuroprotective by transitioning microglia away from the intermediate pro-inflammatory rod-like state, which is aligned with the previously reported anti-inflammatory effects of both Az and Epo. In regions where neuroprotection was seen after Az and Az*Epo combination treatment, there was also increased circularity. While circularity is traditionally thought to be a marker of pro-inflammatory microglia, other signatures of a pro-inflammatory microglial state were absent. This suggests that greater circularity may be associated with multiple microglial phenotypes with different functionalities including secretion of both pro-inflammatory and injury-promoting mediators as well as those that positively affect repair and remyelination (*56, 57*).

By comparison, in regions such as the thalamus where Az and Az*Epo were not neuroprotective, an increased distribution of ameboid and rounded-shaped microglia (SM5) with less cell area coverage was seen compared to OGD. This morphology is suggestive of reactivity involving retraction of cellular processes, swelling of the soma and expression of inflammatory surface antigens when extensive damage and cellular debris is present (*53*). With graphical network analyses, we were able to identify a connection between a greater proportion of microglial with SM5 morphology and increased expression of transcriptional markers of microglial activation and inflammation, including CXCL10, STAT1, and CD86. CXCL10 is regarded as an important mediator in migration and a proinflammatory microglia phenotype as well as initiation of neuroinflammatory processes associated with oligodendrocyte injury (*58*). STAT1 has been linked to microglial response to hypoxia, with CD86 being a common surface marker of pro-inflammatory microglia phenotypes (*59*). As the proportion of SM5 microglia was not increased by OGD alone, it is possible that Az has a direct negative effect on microglia in certain regions that prevents local neuroprotection, a finding that warrants further investigation *in vivo*.

The results of our examination of treatment- and combination-specific neuroprotective mechanisms of Az and Epo suggested emergent responses that were synergistically regulated by Az*Epo, including several pathways associated with cell death and inflammatory pathways including HMGB-1, NLRP3, IRF8, TLR4, CASP3 and PARP1. Though the analysis of pyknotic nuclei cannot necessarily distinguish between cells undergoing different types of programmed cell death, there was some evidence that the combination of Az*Epo had anti-apoptotic effects. For example, in the cortex, Az*Epo triggered the downregulation of Caspase-3, an important executioner caspase that activates nuclear and cytoskeletal protein changes in apoptotic cells (*60*). Through GO analysis, we further inferred that mechanisms of neuroprotection from combinatorial treatment may involve mitigation of oxidative stress in addition to suppression of cellular apoptosis. Investigation of synergistic effects on gene expression also revealed suppression of NLRP3 by Az*Epo, suggesting additional possible effects on cell death via pyroptosis.

We also discovered that the pattern of synergistic neuroprotection by Az*Epo involved both augmentation and reversal of specific transcripts dysregulated by injury. In contrast to the common belief that treatments are only effective when the damage is reversed, we found that augmentation of neurogenic and immunomodulatory responses to OGD was correlated with Az*Epo neuroprotection, potentially due to upregulation of normal reparative responses to injury. Augmenting immune responses to ameliorate disease outcomes is evident elsewhere in the literature. For example, studies exploring the involvement of Th2 cytokine response in pathogenesis of inflammation-induced brain injury have shown that reagents that augment Th2 response improved injury progression (*61, 62*). On the other hand, reversal of other responses to OGD related to oxidative stress, apoptotic cell death, and certain glial differentiation and inflammation-related transcripts were also associated with combinatorial neuroprotection.

Lastly, regional transcriptional state changes were used to compare overlapping and discrete transcriptional responses by group. Despite minimal frank cellular death in the white matter, the greatest number of differentially regulated genes were identified in the white matter, including a greater number that were specifically targeted by Az*Epo. These data suggest that Az*Epo has the potential to result in significant inflammation-meditated neuroprotection in the white matter despite the minimal changes in pyknotic cell counts. While this would need to be confirmed *in vivo*, we speculate that this effect may be more pronounced in a live animal injury model due to the enrichment of transcripts associated with the innate immune response, which is known to play a major role in preterm brain injury (*63*).

The emergent transcriptomic signatures resulting from Az*Epo in combination showcases the advantages of combination therapies by addressing multiple injury mechanisms, increasing the potency and efficacy of treatment. Combinatorial drug therapy can potentially leverage the development of otherwise ineffective monotherapies, allow for repurposing of clinically-approved agents, and accelerate the discovery and development of new therapeutic options (*49*). However, not all effects of Az*Epo could be accurately predicted from their individual treatment responses, suggesting that combinations may need to be screened *de novo*. Future work will investigate the synergistic mechanisms of drug combinations in the OWH ferret slice model, with subsequent application of the most promising combinations *in vivo*, using our ferret model of inflammation-sensitised preterm brain injury (*64*). Clinical translation of preclinical success in combination therapy studies has long presented a challenge in which variables such as dose and administration timing alter therapeutic outcomes. With the help of statistical and computational modelling and screening platforms such as the OWH slice culture model, the efficacy profile of various drug combinations can be screened in a high-throughput manner for reliable determination of additivity or synergism to be clinically translatable.

This study does have several limitations. The hippocampal region in animals of this age is small and we were unable to provide enough processable Iba-1-stained images in the hippocampal region for regional microglia morphological analysis in that region separately. For the same reason, we combined deep grey matter regions for nCounter analysis and therefore did the same for neuroprotection and VAMPIRE analyses to align them. However, some evidence of variability in response to Az*Epo was seen across those regions even though global injury was reduced based on multiple measures. It would be ideal to present and analyse transcriptomic data for all deep grey regions (basal ganglia, thalamus and hippocampus) separately, but this was not technically feasible in a reproducible manner. Additionally, the cell count, VAMPIRE, and transcriptomic data all came from different slices and were aligned by group to perform the analyses, which required us to assume that those responses would be consistent across those slices. However, our group sizes and number of replicates were designed to offset this limitation, and consistent responses were evident (e.g., SM shifts in both Az and Az*Epo groups with corresponding transcriptomic shifts) that suggest that our analyses were robust. The study is also limited by the lack of confirmation of the top transcriptomic responses via proteomic analyses. Advanced proteomic methods in ferret tissue are still in development (*65*), and to our knowledge have never been successfully performed with ferret brain tissue. In addition, relatively few antibodies have been validated for protein quantification in ferret tissue, with the majority focused on models of respiratory disease (*66*). Quantification of protein targets in the model will be the focus of future work.

We also did not have the statistical power to explore sex differences reacting to treatments in regions, though all studies were performed in a sex-balanced manner. As there are sex-dependent differences in cell-specific transcriptomes (*67*), susceptibility to preterm brain damage (*68*), as well as pathophysiological and inflammatory responses (*69*), the exploration of sex-specific injury and treatment mechanisms will be helpful in future studies using this platform. We did not observe statistically significant cell death in the white matter in the extremely preterm-equivalent model, unlike our previously published term-equivalent model (*17*). In order to capture the full spectrum of severity and mechanisms of WMI that disrupts the progression of developmental myelination in preterm infants (*11*), further development of the model may be needed. However, more subtle changes in white matter function as a result of preterm birth and preterm brain injury are common, suggesting that the current model remains highly translation. More broadly, while the OWH model maintains 3D brain structure and function and mimics many regional responses to *in vivo* injury and treatment, it does not incorporate peripheral contributors to brain injury and removes the blood-brain barrier as a regulator of drug uptake into the brain. Therefore, specific pharmacokinetic and *in* vivo efficacy studies of drug combinations will need to be performed as the next stage in translation.

## Conclusions

We established an extremely preterm-equivalent ferret OWH slice model platform to explore regional cellular changes, microglial phenotypic shifts, and spatial transcriptomic signatures in response to injury and treatments. This platform provides a powerful tool to examine combinations of therapeutics to assess potential therapies for preterm brain injury in the gyrencephalic brain. Using this model, we found evidence for synergistic neuroprotection by Az*Epo across the brain, including reversal of multiple inflammatory signatures and associated shifts in microglial morphology. As both Az and Epo are used in EP infants currently and have known safety profiles in that patient population, Az*Epo combination therapy warrants assessment in *in vivo* preterm injury models prior to clinical trials if successful.

## Acknowledgments

The authors would like to thank Mike McKenna and Olivia Dotson for imaging assistance, Olivia White and Lily Farid for their assistance with cell counting, as well as Dr. Katherine Prater for her assistance building the nCounter gene set.

## Author contributions

Conceptualization: TRW, EN, SEJ

Methodology: ZRJ, KAC, OCB, MM, HH, DHM, PB, TRW, EN

Investigation: ZRJ, KAC, OCB, HH, DHM, RB, SEK, TRW

Visualization: ZRJ, OCB, HH, TRW

Funding acquisition: EN, TRW

Project administration: EN, TRW

Supervision: EN, TRW

Writing – original draft: ZRJ, EN, TRW

Writing – review & editing: ZRJ, KAC, OCB, MM, HH, DHM, RB, SEK, PB, SEJ, EN, TRW

## Data, code, and materials availability

All data are available in the main text or the supplementary materials. Code for automated cell counts is available at https://doi.org/10.5061/dryad.r4xgxd2qv.

## List of Supplemental Materials

Figure S1: Experimental schematic of *ex vivo* ferret brain slice oxygen glucose deprivation (OGD) injury model.

Figure S2: Cell death percentage by treatment groups and synergism of Az*Epo combinatorial treatment in the deep grey matter regions.

Figure S3: Regional oligodendrocyte progenitor cell number response to OGD and treatments.

Figure S4: Microglial morphological parameter trends in relation to shape modes.

Figure S5: Principal component analysis plot of nCounter transcriptomics data.

Figure S6: nCounter transcript principal components (PCs) in relation to microglial shape modes and selective morphological parameters.

Figure S7: Spatial transcriptomics panel grouped by pathway categories.

Figure S8: Emergent transcriptomic signatures of synergistic protection by Az*Epo combination treatment by regions.

Table S1: Definitions of microglial morphological parameters

Table S2: Contributions and loadings of top 25 transcripts in all seven nCounter Principal Components (PCs).

Table S3: Top contributing genes in each Principal Component (PC) cluster with log2 fold change and p-value compared to OGD.

Table S4: Differentially expressed genes between OGD and treatments globally, and in the white matter, deep grey matter, and cortex.

## References

1. D. Mota-Rojas et al., Pathophysiology of Perinatal Asphyxia in Humans and Animal Models. Biomedicines 10, 347 (2022).

2. J. C. Y. Lai et al., Immune responses in perinatal brain injury. Brain, Behavior, and Immunity 63, 210–223 (2017).

3. S. Johnson et al., Neurodevelopmental disability through 11 years of age in children born before 26 weeks of gestation. Pediatrics 124, e249–257 (2009).

4. N. Marlow, E. M. Hennessy, M. A. Bracewell, D. Wolke, E. P. S. Group, Motor and executive function at 6 years of age after extremely preterm birth. Pediatrics 120, 793–804 (2007).

5. K. C. K. Kuban et al., Girls and Boys Born before 28 Weeks Gestation: Risks of Cognitive, Behavioral, and Neurologic Outcomes at Age 10 Years. The Journal of Pediatrics 173, 69–75.e61 (2016).

6. S. E. Juul et al., Deaths in a Modern Cohort of Extremely Preterm Infants From the Preterm Erythropoietin Neuroprotection Trial. JAMA Netw Open 5, e2146404 (2022).

7. C. Crump, J. Sundquist, M. A. Winkleby, K. Sundquist, Preterm birth and risk of chronic kidney disease from childhood into mid-adulthood: national cohort study. BMJ 365, l1346 (2019).

8. T. N. K. Raju et al., Long-Term Healthcare Outcomes of Preterm Birth: An Executive Summary of a Conference Sponsored by the National Institutes of Health. The Journal of Pediatrics 181, 309–318.e301 (2017).

9. D. Ely, A. Driscoll, H.-C. Shin, T.-C. Chen, “Infant Mortality by Selected Maternal Characteristics and Race and Hispanic Origin in the United States, 2019-2021,” (2024).

10. J. M. Perlman, White matter injury in the preterm infant: an important determination of abnormal neurodevelopment outcome. Early Human Development 53, 99–120 (1998).

11. S. A. Back, White matter injury in the preterm infant: pathology and mechanisms. Acta Neuropathologica 134, 331–349 (2017).

12. C. Papini et al., Altered Cortical Gyrification in Adults Who Were Born Very Preterm and Its Associations With Cognition and Mental Health. Biological Psychiatry: Cognitive Neuroscience and Neuroimaging 5, 640–650 (2020).

13. T. Wood et al., A Ferret Model of Encephalopathy of Prematurity. Developmental Neuroscience 40, 475–489 (2018).

14. A. R. Barnette et al., Characterization of Brain Development in the Ferret via MRI. Pediatric Research 66, 80–84 (2009).

15. M. D. Agata Tarkowska, Hypoxic-Ischemic Brain Injury after Perinatal Asphyxia as a Possible Factor in the Pathology of Alzheimer’s Disease. Exon Publications, 45–59 (2021).

16. M. M et al., Organotypic whole hemisphere brain slice models to study the effects of donor age and oxygen-glucose-deprivation on the extracellular properties of cortical and striatal tissue. Journal of Biological Engineering 16, (2022).

17. T. R. Wood, et al., A ferret brain slice model of oxygen–glucose deprivation captures regional responses to perinatal injury and treatment associated with specific microglial phenotypes. Bioengineering & Translational Medicine 7, (2022).

18. E. Landucci, D. E. Pellegrini-Giampietro, F. Facchinetti, Experimental Models for Testing the Efficacy of Pharmacological Treatments for Neonatal Hypoxic-Ischemic Encephalopathy. Biomedicines 10, 937 (2022).

19. J. Nair, V. H. S. Kumar, Current and Emerging Therapies in the Management of Hypoxic Ischemic Encephalopathy in Neonates. Children (Basel) 5, 99 (2018).

20. Y. Zhu et al., Identification of novel biomarkers for neonatal hypoxic-ischemic encephalopathy using iTRAQ. Italian Journal of Pediatrics 46, 67 (2020).

21. J. D. E. Barks, Y. Liu, L. Wang, M. P. Pai, F. S. Silverstein, Repurposing Azithromycin for Neonatal Neuroprotection. Pediatric research 86, 444–451 (2019).

22. Y. W. Wu et al., Trial of Erythropoietin for Hypoxic–Ischemic Encephalopathy in Newborns. New England Journal of Medicine 387, 148–159 (2022).

23. S. E. Juul, G. C. Pet, Erythropoietin and Neonatal Neuroprotection. Clin Perinatol 42, 469–481 (2015).

24. C. M. Traudt et al., Concurrent erythropoietin and hypothermia treatment improve outcomes in a term nonhuman primate model of perinatal asphyxia. Dev Neurosci 35, 491–503 (2013).

25. S. E. Juul et al., A Randomized Trial of Erythropoietin for Neuroprotection in Preterm Infants. The New England journal of medicine 382, 233 (2020).

26. L. L. Jantzie et al., Infantile Cocktail of Erythropoietin and Melatonin Restores Gait in Adult Rats with Preterm Brain Injury. Dev Neurosci 44, 266–276 (2022).

27. L. L. Jantzie et al., Repetitive Neonatal Erythropoietin and Melatonin Combinatorial Treatment Provides Sustained Repair of Functional Deficits in a Rat Model of Cerebral Palsy. Front Neurol 9, 233 (2018).

28. S. Robinson et al., Extended Combined Neonatal Treatment With Erythropoietin Plus Melatonin Prevents Posthemorrhagic Hydrocephalus of Prematurity in Rats. Front Cell Neurosci 12, 322 (2018).

29. J. D. E. Barks, Y. Liu, I. A. Dopp, F. S. Silverstein, Azithromycin reduces inflammation-amplified hypoxic–ischemic brain injury in neonatal rats. Pediatric Research 92, 415–423 (2022).

30. D. Amantea et al., Azithromycin protects mice against ischemic stroke injury by promoting macrophage transition towards M2 phenotype. Exp Neurol 275 Pt 1, 116–125 (2016).

31. M. Vrančić et al., Azithromycin distinctively modulates classical activation of human monocytes in vitro. Br J Pharmacol 165, 1348–1360 (2012).

32. S. A. Back, Perinatal white matter injury: the changing spectrum of pathology and emerging insights into pathogenetic mechanisms. Ment Retard Dev Disabil Res Rev 12, 129–140 (2006).

33. G. Favrais et al., Systemic inflammation disrupts the developmental program of white matter. Ann Neurol 70, 550–565 (2011).

34. M. McKenna et al., Organotypic whole hemisphere brain slice models to study the effects of donor age and oxygen-glucose-deprivation on the extracellular properties of cortical and striatal tissue. J Biol Eng 16, 14 (2022).

35. T. E. Oliphant, Python for Scientific Computing. Computing in Science and Engg. 9, 10– 20 (2007).

36. S. van der Walt et al., scikit-image: image processing in Python. PeerJ 2, e453 (2014).

37. G. V. Fabian Pedregosa, Alexandre Gramfort, Vincent Michel, Bertrand Thirion, Olivier Grisel, Mathieu Blondel, Peter Prettenhofer, Ron Weiss, Vincent Dubourg, Jake Vanderplas, Alexandre Passos, David Cournapeau, Matthieu Brucher, Matthieu Perrot, Édouard Duchesnay, Scikit-learn: Machine Learning in Python. Journal of Machine Learning Research 12, 2825–2830 (2011).

38. S. X. Ge, D. Jung, R. Yao, ShinyGO: a graphical gene-set enrichment tool for animals and plants. Bioinformatics 36, 2628–2629 (2020).

39. P. Shannon et al., Cytoscape: A Software Environment for Integrated Models of Biomolecular Interaction Networks. Genome Res 13, 2498–2504 (2003).

40. D. Duarte, N. Vale, Evaluation of synergism in drug combinations and reference models for future orientations in oncology. Current Research in Pharmacology and Drug Discovery 3, 100110 (2022).

41. S. E. Juul et al., Predicting 2-year neurodevelopmental outcomes in extremely preterm infants using graphical network and machine learning approaches. eClinicalMedicine 56, 101782 (2022).

42. D. R. Williams, P. Rast, Back to the basics: Rethinking partial correlation network methodology. Br J Math Stat Psychol 73, 187–212 (2020).

43. S. E. Juul et al., Predicting 2-year neurodevelopmental outcomes in extremely preterm infants using graphical network and machine learning approaches. eClinicalMedicine 56, 101782 (2023).

44. C. Brégère, B. Schwendele, B. Radanovic, R. Guzman, Microglia and Stem-Cell Mediated Neuroprotection after Neonatal Hypoxia-Ischemia. Stem Cell Rev Rep 18, 474–522 (2022).

45. U. Fisch, C. Brégère, F. Geier, L. Chicha, R. Guzman, Neonatal hypoxia-ischemia in rat elicits a region-specific neurotrophic response in SVZ microglia. Journal of Neuroinflammation 17, 26 (2020).

46. J. J. Sun, T.-W. Huang, J. L. Neul, R. S. Ray, Embryonic hindbrain patterning genes delineate distinct cardio-respiratory and metabolic homeostatic populations in the adult | Scientific Reports. Scientific Reports 7, 9117 (2017).

47. C. I. Bliss, The Toxicity of Poisons Applied Jointly1. Annals of Applied Biology 26, 585–615 (1939).

48. Y. Pan, H. Ren, L. Lan, Y. Li, T. Huang, Review of Predicting Synergistic Drug Combinations. Life 13, 1878 (2023).

49. N. R. Twarog, M. Connelly, A. A. Shelat, A critical evaluation of methods to interpret drug combinations. Scientific Reports 10, 5144 (2020).

50. D. P. Schafer, B. Stevens, Microglia Function in Central Nervous System Development and Plasticity. Cold Spring Harb Perspect Biol 7, a020545 (2015).

51. O. Butovsky, H. L. Weiner, Microglial signatures and their role in health and disease. Nat Rev Neurosci 19, 622–635 (2018).

52. W.-S. Qu et al., Rapidly activated epidermal growth factor receptor mediates lipopolysaccharide-triggered migration of microglia. Neurochemistry International 90, 85–92 (2015).

53. J. C. Savage, M. Carrier, M.-È. Tremblay, Morphology of Microglia Across Contexts of Health and Disease. Methods Mol Biol 2034, 13–26 (2019).

54. O. G. Holloway, A. J. Canty, A. E. King, J. M. Ziebell, Rod microglia and their role in neurological diseases. Seminars in Cell & Developmental Biology 94, 96–103 (2019).

55. K. R. Giordano, C. R. Denman, P. S. Dubisch, M. Akhter, J. Lifshitz, An update on the rod microglia variant in experimental and clinical brain injury and disease. Brain Communications 3, fcaa227 (2021).

56. J. L. Mason, K. Suzuki, D. D. Chaplin, G. K. Matsushima, Interleukin-1β Promotes Repair of the CNS. J. Neurosci. 21, 7046–7052 (2001).

57. J. Lee, G. Hamanaka, E. H. Lo, K. Arai, Heterogeneity of microglia and their differential roles in white matter pathology. CNS Neurosci Ther 25, 1290–1298 (2019).

58. T. Clarner et al., CXCL10 Triggers Early Microglial Activation in the Cuprizone Model. The Journal of Immunology 194, 3400–3413 (2015).

59. E. Butturini, D. Boriero, A. Carcereri de Prati, S. Mariotto, STAT1 drives M1 microglia activation and neuroinflammation under hypoxia. Archives of Biochemistry and Biophysics 669, 22–30 (2019).

60. S. Elmore, Apoptosis: a review of programmed cell death. Toxicol Pathol 35, 495–516 (2007).

61. U. Gimsa, S. A. Wolf, D. Haas, I. Bechmann, R. Nitsch, Th2 cells support intrinsic anti-inflammatory properties of the brain. Journal of Neuroimmunology 119, 73–80 (2001).

62. P. L. Vieira, H. C. Heystek, J. Wormmeester, E. A. Wierenga, M. L. Kapsenberg, Glatiramer Acetate (Copolymer-1, Copaxone) Promotes Th2 Cell Development and Increased IL-10 Production Through Modulation of Dendritic Cells1. The Journal of Immunology 170, 4483–4488 (2003).

63. J. Herz, I. Bendix, U. Felderhoff-Muser, Peripheral immune cells and perinatal brain injury: a double-edged sword? Pediatr Res 91, 392–403 (2022).

64. K. A. Corry et al., Evaluating Neuroprotective Effects of Uridine, Erythropoietin, and Therapeutic Hypothermia in a Ferret Model of Inflammation-Sensitized Hypoxic-Ischemic Encephalopathy. International Journal of Molecular Sciences 22, 9841 (2021).

65. N. Ali et al., Semen proteome and transcriptome of the endangered black-footed ferret (Mustela nigripes) show association with the environment and fertility outcome. Sci Rep 14, 7063 (2024).

66. N. He, et al., Ferret models of alpha-1 antitrypsin deficiency develop lung and liver disease. JCI Insight 7, (2022).

67. I. Kremsky et al., Fetal hypoxia results in sex- and cell type-specific alterations in neonatal transcription in rat oligodendrocyte precursor cells, microglia, neurons, and oligodendrocytes. Cell Biosci 13, 58 (2023).

68. S. R. Mayoral, G. Omar, A. A. Penn, Sex Differences in a Hypoxia Model of Preterm Brain Damage. Pediatric Research 66, 248–253 (2009).

69. L. Beckmann et al., Regulatory T Cells Contribute to Sexual Dimorphism in Neonatal Hypoxic-Ischemic Brain Injury. Stroke 53, 381–390 (2022).

