## Supplemental Figures for "Multi-modal screening for synergistic neuroprotection of extremely preterm brain injury"

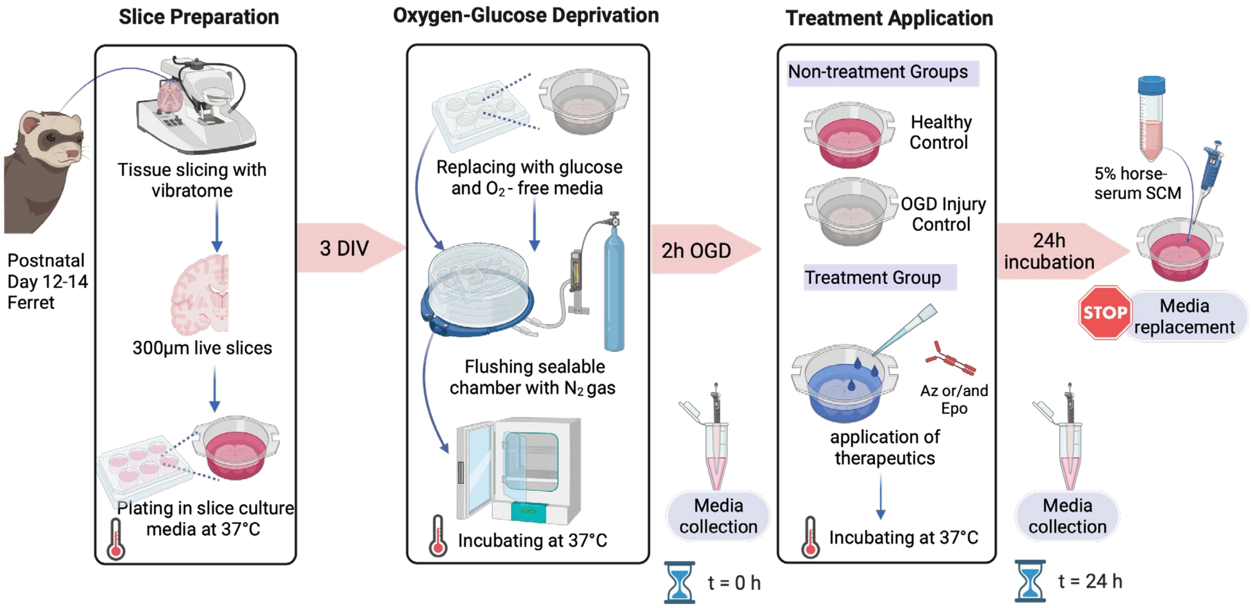


**Figure S1. Experimental schematic of *ex vivo* ferret brain slice oxygen glucose deprivation (OGD) injury model.** 300 µm thick OWH slices were harvested with a vibratome in dissection media from P12-14 ferrets. Slices were cultured for 3 days *in vitro* (DIV), then OGD was applied by replacing standard culture media (SCM) with OGD media and placing slices inside a nitrogen-purged hypoxic (0% $O_2$) chamber at 37°C for two hours. Afterwards, OWH slices were returned to normoxia conditions (5% $CO_2$, balance air) at 37°C in SCM. Treatments were applied topically immediately after OGD in SCM for 24h. Post-OGD media and post-treatment media were collected for further assessment.

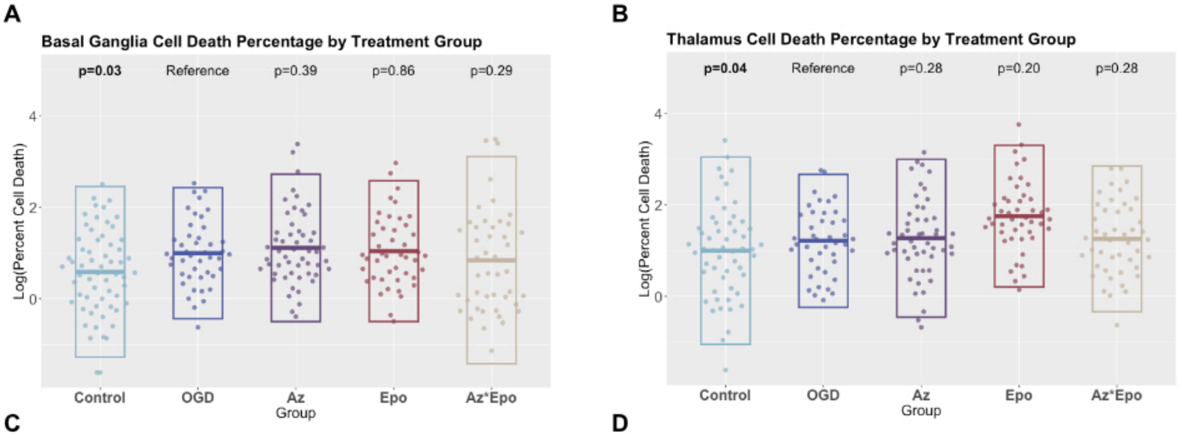


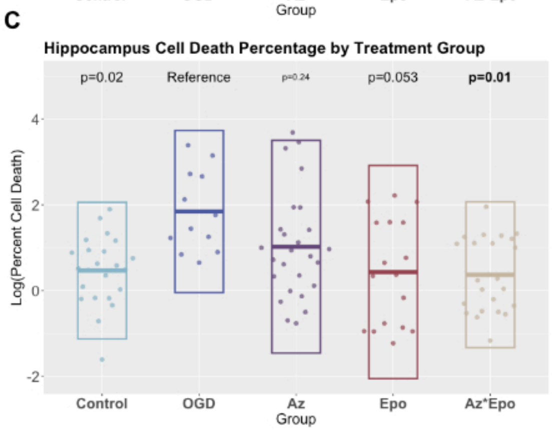


D

| | Deep Grey Matter Regions | | |
|---|---|---|---|
| **Model** | **BG** | **Thal** | **Hipp** |
| Combination Subthresholding | x | x | ✓ |
| Highest Single Agent | ✓ | ✓ | ✓ |
| Response Additivity | ✓ | ✓ | x |
| Bliss Independence | NA (✓) | NA (✓) | ✓ |

**Figure S2. Cell death percentage by treatment groups and synergism of Az*Epo combinatorial treatment in the deep grey matter regions.** Log-transformed cell death percentage by treatment groups in the deep grey regions (A) basal ganglia, (B) thalamus, and (C) hippocampus are shown. Each point represents one of n=3-4 individual images per region per group from n=9 slices. Boxes show mean with SD. Bolded p-values indicate significant differences compared to the untreated OGD group performed with linear mixed effects models with fixed effects of region and random effect of slice. (D) Synergism of Az*Epo combinatorial treatment evaluated by regional cell death percentage in the deep grey matter regions compared to Az, Epo individual treatments. Green ticks indicate evidence for synergy under that model in that region/assessemnt type. Purple x's are displayed when no evidence of synergy is seen. NA with a tick indicates where the conditions of Bliss Independence were not fully met (one treatment did not display any effect on its own) but synergy was suggested according to the model.

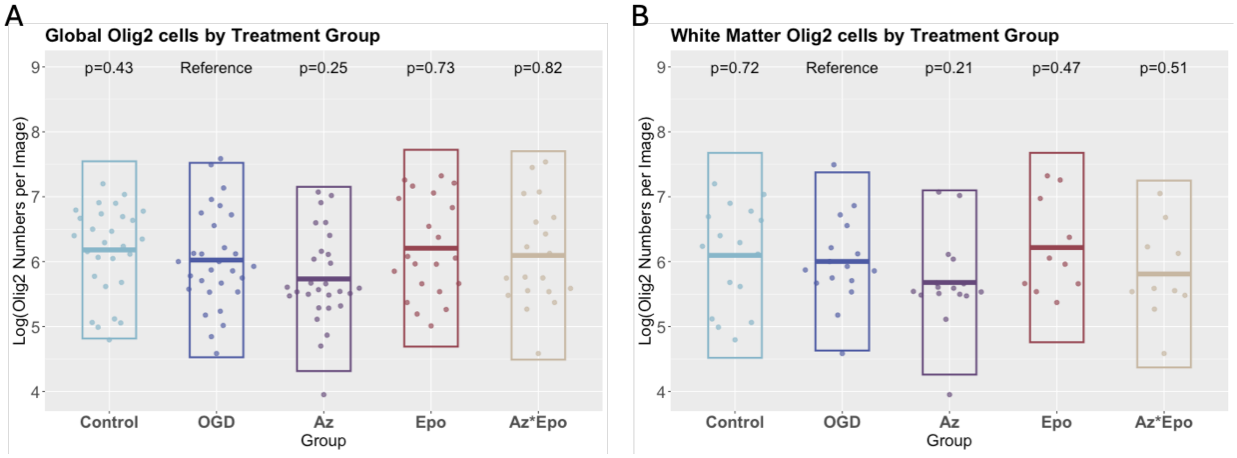


**Figure S3. Regional oligodendrocyte progenitor cell number response to OGD and treatments.** Olig2+ cell counts by treatment groups (A) globally (all regions) and (B) in the subcortical white matter. Each point represents one of n = 3-4 individual images per region per group from n = 9 slices. Boxes show mean with SD. Bolded p-values indicate significant differences compared to the untreated OGD group performed with linear mixed effects models with fixed effects of region and random effect of slice.

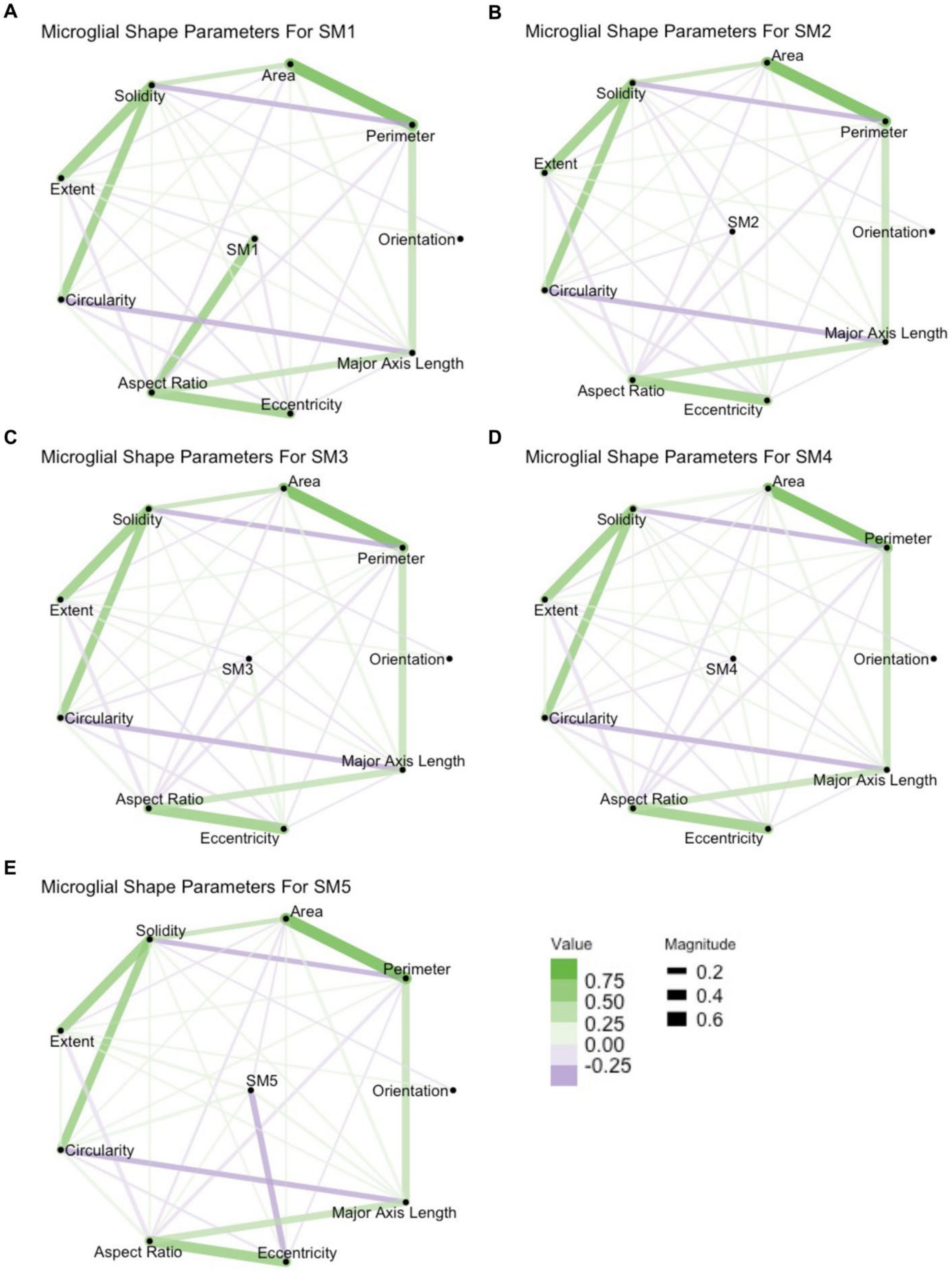

A
Microglial Shape Parameters For SM1
Area
Solidity
Perimeter
Extent
SM1
Orientation
Circularity
Major Axis Length
Aspect Ratio
Eccentricity
B
Microglial Shape Parameters For SM2
Area
Solidity
Perimeter
Extent
SM2
Orientation
Circularity
Major Axis Length
Aspect Ratio
Eccentricity
C
Microglial Shape Parameters For SM3
Area
Solidity
Perimeter
Extent
SM3
Orientation
Circularity
Major Axis Length
Aspect Ratio
Eccentricity
D
Microglial Shape Parameters For SM4
Area
Solidity
Perimeter
Extent
SM4
Orientation
Circularity
Major Axis Length
Aspect Ratio
Eccentricity
E
Microglial Shape Parameters For SM5
Solidity
Area
Perimeter
Extent
SM5
Orientation
Circularity
Major Axis Length
Aspect Ratio
Eccentricity
Value
0.75
0.50
0.25
0.00
-0.25
Magnitude
0.2
0.4
0.6

**Figure S4. Microglial morphological parameter trends in relation to shape modes.** Graphical network analyses of relationships between morphological parameters and microglial SMs (A) 1, (B) 2, (C) 3, (D) 4, and (E) 5 are shown. The morphological parameters were quantified from microglial cells captured from n = 3-4 individual images per region from n = 9 slices in each group. Green lines indicate significant positive correlations between the morphological parameters and SMs. Purple lines indicate significant negative correlations between the morphological parameters and SMs. The thickness of lines in graphical networks imply how positively or negatively related the relationships were.

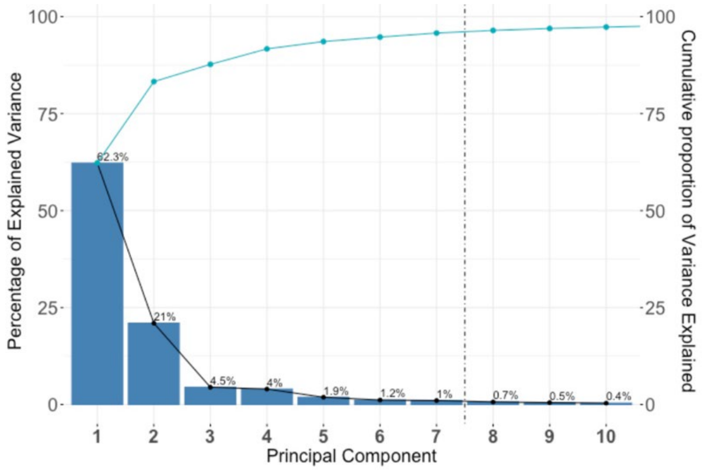


**Figure S5. Principal component analysis plot of nCounter transcriptomics data**. The screen plot shows the separation of different groupings of the normalised transcript levels screened in the NanoString nCounter panel. We selected the first seven PCs that explained at least 95% of the variance of the nCounter data in this study.

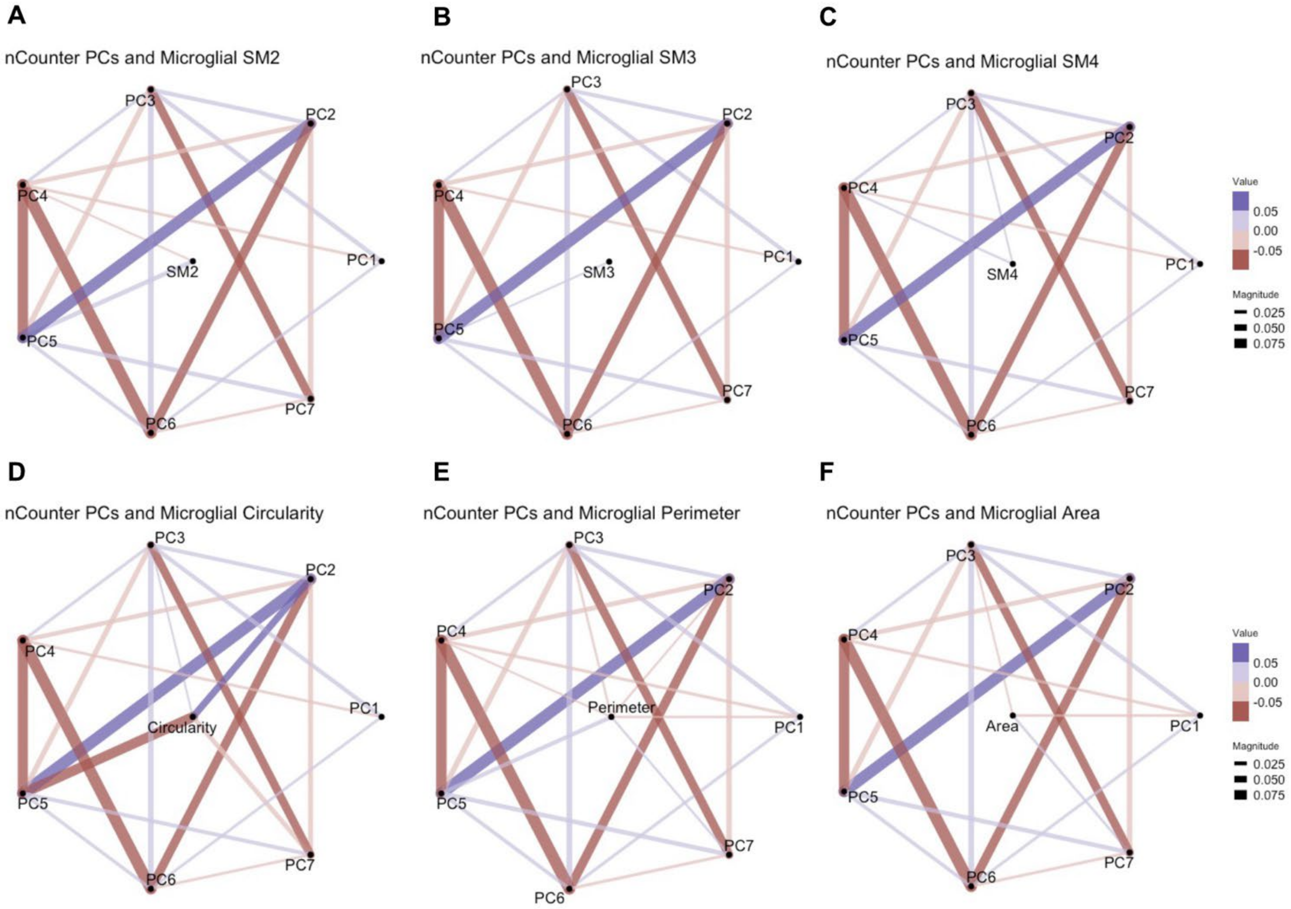


**Figure S6. nCounter transcript principal components (PCs) in relation to microglial shape modes and selective morphological parameters.** Graphical network analyses of seven PCs of nCounter data in relation to microglial SMs (A) 2, (B) 3, and (C) 4 are shown. In addition, networks of nCounter PCs in relation to microglia morphological parameters, including (D) circularity, (E) perimeter, and (F) area are shown. Purple lines indicate significant positive correlation between the nCounter PCs and microglial SMs. Red lines indicate significant negative correlation between nCounter PCs and microglial SMs. The thickness of lines in graphical networks imply how positively or negatively correlated the relationships were.

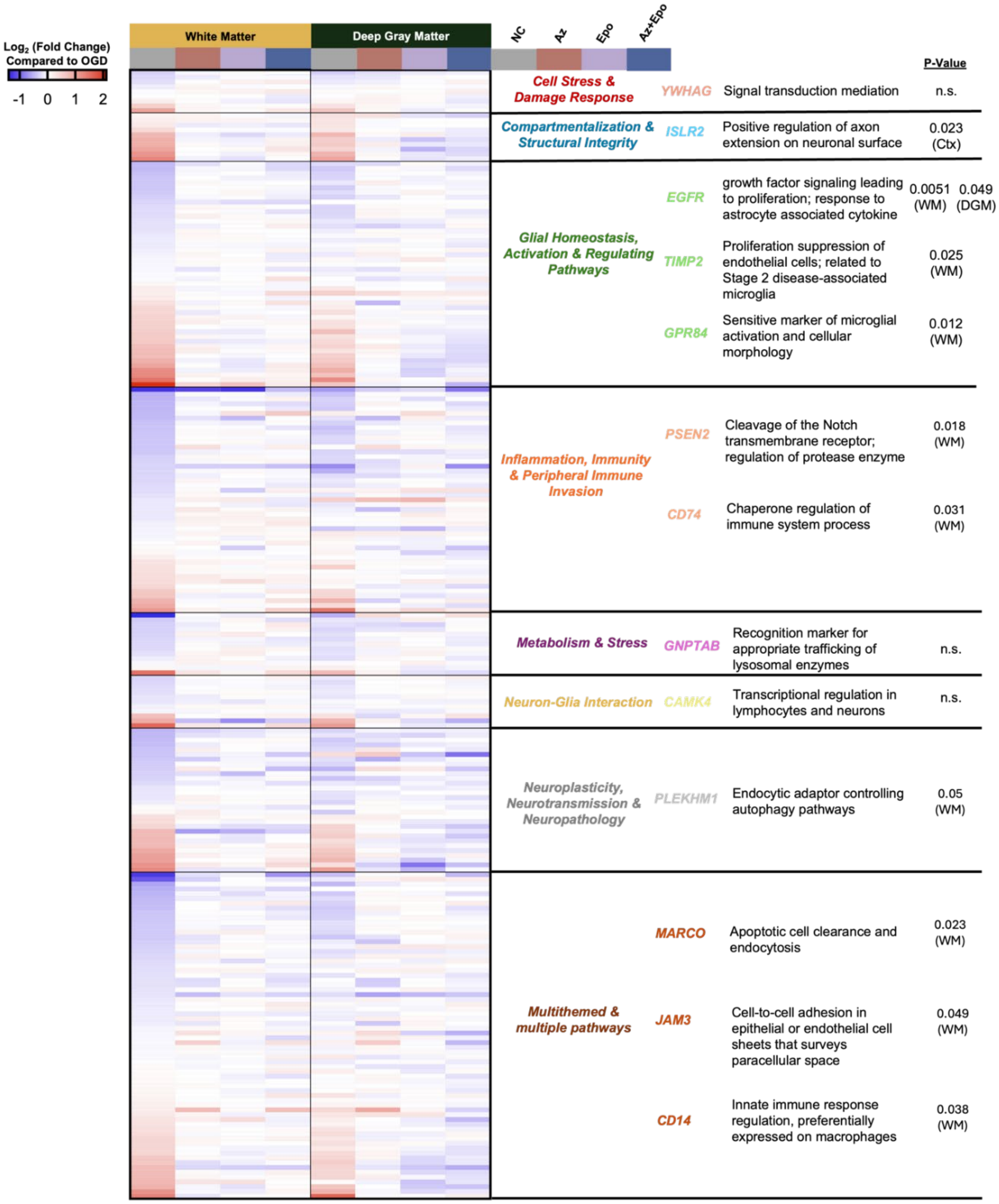


**Figure S7. Spatial transcriptomics panel grouped by pathway categories.** The heatmap shows $Log_2$ (fold change) in the white and deep grey matter for each group compared to OGD. 235 endogenous genes were assessed and categorised into seven different categories by functionality, including a group of multithemed genes that had at least three categorial functions listed. Representative genes in each category that were increased by OGD but regulated by Az*Epo ($p<0{\cdot}05$, t-test) to show similar fold change trends as the NC group are listed. Their respective functions and p-value within the region involved are shown.

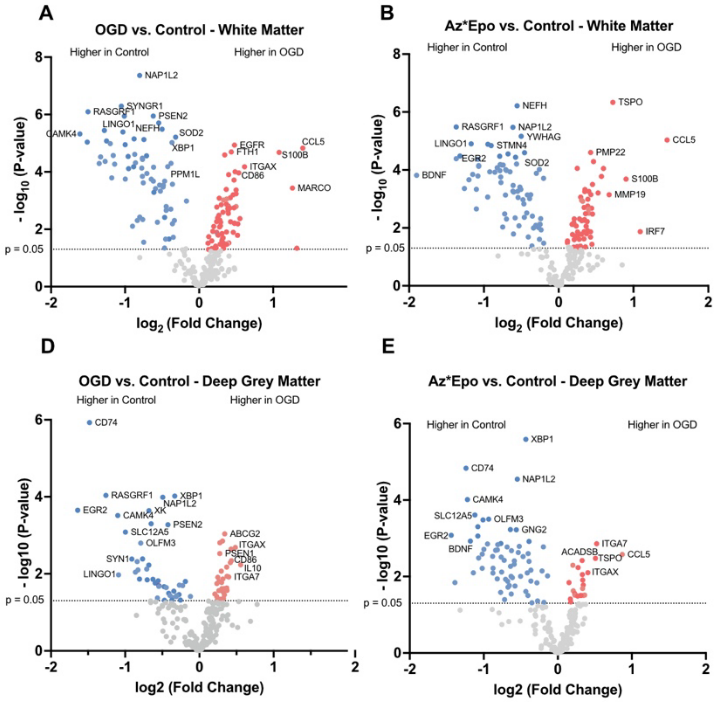


C

| | Log2 Fold-Change (White Matter) | | | |
|---|---|---|---|---|
| **Probe** | **OGD vs. Con** | **Az vs. Con** | **Epo vs. Con** | **Az*Epo vs.Con** |
| **MARCO** | **1.26** | **1.05** | **1.17** | 0.52 |
| **GSN** | **0.50** | **0.42** | **0.30** | 0.25 |
| **PLXNC1** | **0.42** | **0.44** | **0.39** | 0.22 |
| **CNTF** | **0.39** | **0.36** | **0.34** | 0.22 |
| **NLRP3** | **0.33** | **0.41** | **0.41** | 0.31 |
| **GBA** | **0.29** | **0.21** | **0.22** | 0.11 |
| **ABL1** | **0.27** | **0.31** | **0.18** | 0.14 |
| **SCAMP2** | **0.25** | **0.24** | **0.23** | 0.10 |
| **RHOA** | **0.25** | **0.23** | **0.21** | 0.16 |
| **TLR4** | **0.24** | **0.25** | **0.29** | 0.14 |
| **TGFBR1** | **0.17** | **0.30** | **0.21** | 0.14 |
| **C3** | **-0.83** | **-0.87** | **-0.79** | -0.50 |

Key: 1, 0.5, 0, -0.5, -1

F

| | Log2 Fold-Change (Deep Grey Matter) | | | |
|---|---|---|---|---|
| **Probe** | **OGD vs. Con** | **Az vs. Con** | **Epo vs. Con** | **Az*Epo vs.Con** |
| **IL10** | **0.56** | **0.40** | **0.51** | 0.38 |
| **CD86** | **0.43** | **0.48** | **0.48** | 0.28 |
| **PLXNC1** | **0.43** | **0.44** | **0.38** | 0.23 |
| **JAM3** | **0.37** | **0.54** | **0.43** | 0.15 |
| **EGFR** | **0.37** | **0.41** | **0.31** | 0.11 |
| **CASP8** | **0.37** | **0.37** | **0.38** | 0.22 |
| **GSN** | **0.34** | **0.47** | **0.30** | 0.07 |
| **SLC1A3** | **0.30** | **0.35** | **0.24** | 0.24 |
| **FIG4** | **0.28** | **0.38** | **0.33** | 0.12 |
| **ITGB5** | **0.26** | **0.36** | **0.31** | 0.17 |
| **MAPK3** | **0.25** | **0.24** | **0.23** | 0.16 |
| **GNPTAB** | **0.24** | **0.33** | **0.25** | 0.08 |
| **LMNA** | **0.24** | **0.31** | **0.26** | 0.18 |
| **CDS1** | **-0.27** | **-0.39** | **-0.33** | -0.21 |
| **CD14** | **-0.31** | **-0.41** | **-0.30** | -0.17 |
| **IGF1** | **-0.40** | **-0.51** | **-0.51** | -0.30 |

**Figure S8. Emergent transcriptomic signatures of synergistic protection by Az*Epo combination treatment by regions.** Volcano plots of differentially expressed genes after OGD (A) and Az*Epo combination treatment (B) in the white matter, as well as differentially expressed genes after OGD (D) and Az*Epo combination treatment (E) in the deep grey matter. Coloured dots represent genes that were significantly differentially expressed comparing with the control group with p value < 0·05. Positive $\log_2$ fold-change (red) indicates upregulated expression after OGD or Az*Epo combination treatment, and negative $\log_2$ fold-change (blue) indicates downregulated expression. The heatmap shows $\log_2$ fold-change for each group compared to control in the white matter (C) and in the deep grey matter (F). Bolded values are biomarkers that are significantly differentially expressed compared to control (p<0·05, t-test). The genes that were normalised by Az*Epo but not by Az or Epo individually are shown. N = 6 slices per region from each treatment group.

### Table S1. Definitions of microglial morphological parameters.

| | **Morphological Parameters** | **Definition** |
|---|---|---|
| 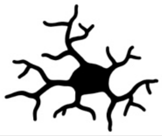 | Area (**A**) | area coverage of the cell body |
| 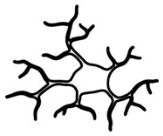 | Perimeter (**P**) | perimeter of the cell body |
| 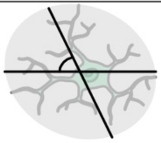 | Orientation | angle between the horizontal line and the major axis of the ellipse that encompasses the cell |
| 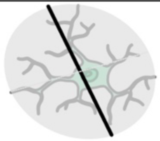 | Major Axis Length | the longest diameter of the ellipse that encompasses the cell |
| 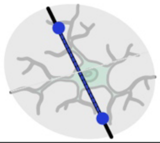 | Eccentricity | ratio of the distance between focal points over the major axis length of the ellipse |
| 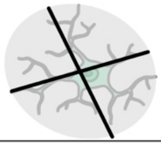 | Aspect Ratio | ratio of the major and minor axis length of the ellipse |
| 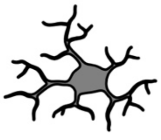 | Circularity (**$4\pi A/P^2$**) | circularity of the cell body |
| 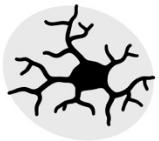 | Extent | ratio of the cell area and the bounding ellipse |
| 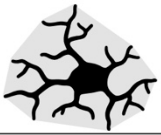 | Solidity | ratio of the cell area and the convex hull area |

### Table S2. Contributions and loadings of top 25 transcripts in all seven nCounter PCs.

| | PC1 | |
|---|---|---|
| | Contribution | Loading |
| IL1B | 0.62767 | 0.07923 |
| HAMP | 0.62767 | 0.07923 |
| MOG | 0.62745 | 0.07921 |
| GDNF | 0.62741 | 0.07921 |
| GPR84 | 0.62739 | 0.07921 |
| KLK6 | 0.62735 | 0.07921 |
| NTRK1 | 0.62731 | 0.07920 |
| IL8 | 0.62730 | 0.07920 |
| NOS2 | 0.62730 | 0.07920 |
| MAG | 0.62729 | 0.07920 |
| BCAS1 | 0.62728 | 0.07920 |
| RET | 0.62723 | 0.07920 |
| UBE4B | 0.62723 | 0.07920 |
| PLXNB3 | 0.62722 | 0.07920 |
| GJB1 | 0.62721 | 0.07920 |
| CSPG4 | 0.62721 | 0.07920 |
| ADRB1 | 0.62720 | 0.07920 |
| NGFR | 0.62719 | 0.07920 |
| NINJ2 | 0.62718 | 0.07919 |
| TREM1 | 0.62715 | 0.07919 |
| IL10 | 0.62714 | 0.07919 |
| IL6 | 0.62714 | 0.07919 |
| TNF | 0.62712 | 0.07919 |
| ABCG2 | 0.62711 | 0.07919 |
| CD300LF | 0.62711 | 0.07919 |

| | PC2 | |
|---|---|---|
| | Contribution | Loading |
| STMN4 | 1.7962 | -0.1340 |
| CDK5R1 | 1.7679 | -0.1330 |
| SYT1 | 1.7482 | -0.1322 |
| THY1 | 1.7050 | -0.1306 |
| YWHAH | 1.6735 | -0.1294 |
| LRRC7 | 1.6696 | -0.1292 |
| DCLK1 | 1.6417 | -0.1281 |
| RTN1 | 1.6297 | -0.1277 |
| STMN1 | 1.6213 | -0.1273 |
| FGF13 | 1.5958 | -0.1263 |
| YWHAG | 1.5582 | -0.1248 |
| SCN2A | 1.5454 | -0.1243 |
| SNCA | 1.4500 | -0.1204 |
| PRKCZ | 1.4473 | -0.1203 |
| GAS6 | 1.4249 | 0.1194 |
| YWHAZ | 1.4246 | -0.1194 |
| SYT4 | 1.4201 | -0.1192 |
| OPCML | 1.4129 | -0.1189 |
| NPTN | 1.4077 | -0.1186 |
| SPTAN1 | 1.3930 | -0.1180 |
| TREM2 | 1.3923 | 0.1180 |
| ITGB5 | 1.3615 | 0.1167 |
| AXL | 1.3168 | 0.1148 |
| MEF2A | 1.2589 | -0.1122 |
| STXBP1 | 1.2513 | -0.1119 |

| | PC3 | |
|---|---|---|
| | Contribution | Loading |
| SOD1 | 5.7355 | 0.2395 |
| MAPK1 | 5.3700 | 0.2317 |
| HDAC2 | 4.9576 | 0.2227 |
| ETV5 | 3.6795 | 0.1918 |
| HIF1A | 3.4227 | 0.1850 |
| ABL1 | 3.3835 | 0.1839 |
| HMGB1 | 3.3073 | 0.1819 |
| CDC42 | 3.1464 | 0.1774 |
| NF1 | 2.4736 | 0.1573 |
| FN1 | 2.4209 | 0.1556 |
| FIG4 | 2.4091 | 0.1552 |
| ATP6V1B2 | 2.2852 | 0.1512 |
| MAPK9 | 2.0544 | 0.1433 |
| LMNA | 2.0489 | 0.1431 |
| PSEN1 | 1.9375 | 0.1392 |
| RHOA | 1.9166 | 0.1384 |
| NCKAP1 | 1.8620 | 0.1365 |
| GSN | 1.8056 | 0.1344 |
| FTH1 | 1.7934 | 0.1339 |
| SPP1 | 1.7501 | 0.1323 |
| SLC1A3 | 1.7405 | 0.1319 |
| LGALS3 | 1.6606 | 0.1289 |
| ACSL4 | 1.6521 | 0.1285 |
| JAM3 | 1.4988 | 0.1224 |
| MBP | 1.3523 | -0.1163 |

| | PC4 | |
|---|---|---|
| | Contribution | Loading |
| CCL5 | 6.8727 | 0.2622 |
| MMP12 | 4.4199 | 0.2102 |
| FN1 | 4.4150 | 0.2101 |
| SPP1 | 4.0714 | 0.2018 |
| PPM1L | 3.7162 | -0.1928 |
| ATP6V1B2 | 3.2150 | -0.1793 |
| LMNA | 2.7742 | 0.1666 |
| PLP1 | 2.6978 | -0.1643 |
| SIRT2 | 2.4793 | -0.1575 |
| OLIG2 | 2.4267 | -0.1558 |
| ENPP6 | 2.4046 | -0.1551 |
| COL6A3 | 2.3784 | 0.1542 |
| PIK3R1 | 2.2871 | -0.1512 |
| MBP | 2.2832 | -0.1511 |
| C3 | 2.1943 | -0.1481 |
| CSF1R | 1.6660 | -0.1291 |
| MAPK9 | 1.6348 | -0.1279 |
| LGALS3 | 1.6274 | 0.1276 |
| GSN | 1.6001 | 0.1265 |
| STAT1 | 1.4978 | -0.1224 |
| MYRF | 1.4069 | -0.1186 |
| P2RY12 | 1.3827 | -0.1176 |
| CLDN5 | 1.3473 | 0.1161 |
| TIMP2 | 1.3256 | 0.1151 |
| SOX10 | 1.2662 | -0.1125 |

| | PC5 | |
|---|---|---|
| | Contribution | Loading |
| SERPINE2 | 12.2595 | 0.3501 |
| CAMK2G | 10.5489 | 0.3248 |
| S100B | 7.8118 | 0.2795 |
| MMP2 | 7.7825 | 0.2790 |
| CXCL10 | 6.6603 | -0.2581 |
| CLDN5 | 3.9668 | 0.1992 |
| STAT1 | 3.2028 | -0.1790 |
| PIK3R1 | 2.6759 | 0.1636 |
| GAS6 | 2.1551 | 0.1468 |
| SLC1A3 | 1.7489 | 0.1322 |
| ITGA7 | 1.5733 | 0.1254 |
| FN1 | 1.5120 | 0.1230 |
| ETV5 | 1.4889 | -0.1220 |
| CD86 | 1.4366 | -0.1199 |
| SOD1 | 1.3385 | 0.1157 |
| COL6A3 | 1.3239 | 0.1151 |
| MBP | 1.1496 | 0.1072 |
| CD9 | 1.1255 | 0.1061 |
| C3 | 1.1209 | 0.1059 |
| CTSB | 1.0624 | -0.1031 |
| PSMB9 | 0.9991 | -0.1000 |
| EGFR | 0.9825 | -0.0991 |
| CYBB | 0.9804 | -0.0990 |
| TIMP2 | 0.9792 | 0.0990 |
| PSMB8 | 0.9524 | -0.0976 |

| | PC6 | |
|---|---|---|
| | Contribution | Loading |
| GLUL | 16.5381 | 0.4067 |
| LGALS3 | 5.8749 | -0.2424 |
| CTSB | 5.5475 | 0.2355 |
| GPR34 | 4.1485 | -0.2037 |
| CLDN5 | 3.7744 | 0.1943 |
| STAT1 | 3.4281 | 0.1852 |
| MMP14 | 3.4094 | -0.1846 |
| TYROBP | 2.6155 | 0.1617 |
| C3 | 2.3194 | -0.1523 |
| PRKAG1 | 2.0895 | -0.1446 |
| SPP1 | 2.0719 | -0.1439 |
| PLA2G4A | 2.0212 | -0.1422 |
| PSMB9 | 1.6631 | 0.1290 |
| CSF1R | 1.5763 | 0.1255 |
| ITGAX | 1.4378 | 0.1199 |
| MBP | 1.2820 | -0.1132 |
| HDAC2 | 1.2695 | -0.1127 |
| CXCL10 | 1.2278 | 0.1108 |
| PSMB8 | 1.2237 | 0.1106 |
| CDC42 | 1.2213 | -0.1105 |
| ENPP6 | 1.0923 | -0.1045 |
| CLEC7A | 1.0517 | 0.1026 |
| UGT8 | 0.8692 | -0.0932 |
| OLIG2 | 0.8626 | 0.0929 |
| S100B | 0.8453 | 0.0919 |

| | PC7 | |
|---|---|---|
| | Contribution | Loading |
| SOX2 | 7.6021 | -0.2757 |
| DLX2 | 7.4996 | -0.2739 |
| ABL1 | 5.3898 | -0.2322 |
| C3 | 5.0020 | 0.2237 |
| SOX9 | 4.1339 | -0.2033 |
| TSC1 | 3.9322 | -0.1983 |
| SPP1 | 3.8858 | 0.1971 |
| TYROBP | 3.2553 | 0.1804 |
| CTSD | 2.9986 | 0.1732 |
| CTSB | 2.7522 | 0.1659 |
| PRKAG1 | 2.6622 | -0.1632 |
| GSN | 2.0964 | -0.1448 |
| DLX1 | 1.8885 | -0.1374 |
| ATP6V1B2 | 1.8884 | 0.1374 |
| CXCL10 | 1.6944 | 0.1302 |
| ACSL4 | 1.6130 | 0.1270 |
| CCL5 | 1.4460 | 0.1203 |
| LGALS3 | 1.3292 | 0.1153 |
| IGF1 | 1.2096 | 0.1100 |
| SLC1A3 | 1.2035 | -0.1097 |
| NF1 | 1.1715 | -0.1082 |
| ADRA2A | 1.1149 | -0.1056 |
| CYBB | 1.0430 | 0.1021 |
| SPI1 | 0.9979 | 0.0999 |
| PPM1L | 0.9607 | -0.0980 |

| Contribution |
|---|
| 16 |
| 12 |
| 8 |
| 4 |
| 0 |

| Loading |
|---|
| 0.4 |
| 0.2 |
| 0 |
| -0.2 |
| -0.4 |

**Table S3. Top contributing genes in each PC cluster with log2 fold change and p-value compared to OGD.**

| | PC1 | | | | | |
|---|---|---|---|---|---|---|
| | Az | | Epo | | Az+Epo | |
| | Log2 fold change | P-value | Log2 fold change | P-value | Log2 fold change | P-value |
| GPR84 | 0.409 | 0.008 | 0.232 | 0.125 | 0.489 | 0.001 |
| IL8 | 0.150 | 0.447 | -0.096 | 0.627 | -0.054 | 0.785 |
| MAG | -3.600 | 2.010E-11 | -3.270 | 5.060E-10 | -3.490 | 6.180E-11 |
| RET | 0.013 | 0.937 | 0.023 | 0.886 | 0.036 | 0.819 |
| UBE4B | 0.004 | 0.943 | -0.098 | 0.064 | -0.149 | 0.005 |
| PLXNB3 | -0.231 | 0.205 | -0.182 | 0.316 | -0.067 | 0.712 |
| CSPG4 | 0.095 | 0.422 | 0.086 | 0.467 | 0.018 | 0.878 |
| NGFR | -0.260 | 0.095 | -0.210 | 0.176 | -0.134 | 0.386 |
| TREM1 | 0.026 | 0.866 | 0.065 | 0.673 | 0.167 | 0.280 |
| IL10 | 0.006 | 0.964 | 0.027 | 0.852 | -0.060 | 0.674 |
| IL6 | -0.116 | 0.545 | -0.074 | 0.698 | -0.045 | 0.814 |
| ABCG2 | -0.192 | 0.047 | -0.165 | 0.087 | -0.244 | 0.012 |
| CD300LF | 0.153 | 0.402 | 0.041 | 0.823 | 0.113 | 0.536 |
| MOBP | 0.124 | 0.486 | 0.059 | 0.742 | 0.027 | 0.880 |
| MAFB | -4.500 | 5.170E-15 | -4.490 | 5.940E-15 | -4.370 | 1.900E-14 |

| | PC2 | | | | | |
|---|---|---|---|---|---|---|
| | Az | | Epo | | Az+Epo | |
| | Log2 fold change | P-value | Log2 fold change | P-value | Log2 fold change | P-value |
| YWHAG | 0.963 | 0.0627 | 0.833 | 0.107 | 0.847 | 0.101 |
| DCLK1 | 0.798 | 0.0865 | 0.627 | 0.176 | 0.713 | 0.125 |
| SNCA | 0.378 | 0.147 | 0.0669 | 0.797 | 0.338 | 0.195 |
| YWHAH | 0.239 | 0.308 | 0.0951 | 0.684 | 0.328 | 0.163 |
| SCN2A | 0.211 | 0.454 | 0.0392 | 0.889 | 0.268 | 0.341 |
| CDK5R1 | 0.295 | 0.232 | 0.0913 | 0.711 | 0.233 | 0.345 |
| SYT1 | 0.292 | 0.312 | 0.076 | 0.792 | 0.229 | 0.426 |
| STMN1 | 0.184 | 0.45 | 0.0166 | 0.946 | 0.198 | 0.415 |
| PRKCZ | 0.202 | 0.363 | 0.0903 | 0.684 | 0.157 | 0.48 |
| LRRC7 | 0.327 | 0.175 | 0.0713 | 0.766 | 0.148 | 0.538 |
| STMN4 | 0.125 | 0.64 | -0.0399 | 0.881 | 0.0982 | 0.712 |
| FGF13 | 0.245 | 0.357 | 0.0607 | 0.819 | 0.0964 | 0.716 |
| RTN1 | 0.256 | 0.327 | 0.00663 | 0.98 | 0.0877 | 0.736 |
| THY1 | 0.136 | 0.579 | 0.0472 | 0.848 | 0.0807 | 0.742 |
| GAS6 | -0.0303 | 0.885 | 0.0649 | 0.758 | 0.0609 | 0.772 |

| | PC3 | | | | | |
|---|---|---|---|---|---|---|
| | Az | | Epo | | Az+Epo | |
| | Log2 fold change | P-value | Log2 fold change | P-value | Log2 fold change | P-value |
| CDC42 | 0.89 | 0.081 | 0.85 | 0.0952 | 0.736 | 0.148 |
| ATP6V1B2 | 0.757 | 0.123 | 0.757 | 0.123 | 0.665 | 0.175 |
| LMNA | 0.693 | 0.119 | 0.648 | 0.145 | 0.618 | 0.164 |
| SOD1 | 0.691 | 0.139 | 0.703 | 0.132 | 0.596 | 0.201 |
| MAPK1 | 0.786 | 0.0896 | 0.694 | 0.133 | 0.585 | 0.204 |
| PSEN1 | 0.759 | 0.0683 | 0.694 | 0.0951 | 0.576 | 0.164 |
| NF1 | 0.832 | 0.068 | 0.732 | 0.108 | 0.574 | 0.206 |
| HIF1A | 0.713 | 0.122 | 0.611 | 0.185 | 0.531 | 0.249 |
| ABL1 | 0.716 | 0.0859 | 0.606 | 0.145 | 0.529 | 0.202 |
| HDAC2 | 0.733 | 0.11 | 0.658 | 0.151 | 0.518 | 0.257 |
| MAPK9 | 0.61 | 0.127 | 0.523 | 0.19 | 0.488 | 0.222 |
| ETV5 | 0.604 | 0.123 | 0.576 | 0.141 | 0.444 | 0.255 |
| FIG4 | 0.641 | 0.105 | 0.571 | 0.148 | 0.398 | 0.312 |
| FN1 | 0.258 | 0.292 | 0.101 | 0.679 | 0.061 | 0.803 |
| HMGB1 | 0.417 | 0.144 | 0.238 | 0.403 | -0.244 | 0.392 |

| | PC4 | | | | | |
|---|---|---|---|---|---|---|
| | Az | | Epo | | Az+Epo | |
| | Log2 fold change | P-value | Log2 fold change | P-value | Log2 fold change | P-value |
| PIK3R1 | 0.721 | 0.116 | 0.735 | 0.109 | 0.703 | 0.125 |
| PPM1L | 0.78 | 0.0551 | 0.666 | 0.101 | 0.689 | 0.0893 |
| ATP6V1B2 | 0.757 | 0.123 | 0.757 | 0.123 | 0.665 | 0.175 |
| SIRT2 | 0.607 | 0.223 | 0.631 | 0.205 | 0.626 | 0.208 |
| LMNA | 0.693 | 0.119 | 0.648 | 0.145 | 0.618 | 0.164 |
| OLIG2 | 0.523 | 0.237 | 0.56 | 0.206 | 0.588 | 0.184 |
| MMP12 | 0.629 | 0.029 | 0.3 | 0.292 | 0.497 | 0.0826 |
| CCL5 | 0.351 | 0.192 | 0.141 | 0.598 | 0.221 | 0.41 |
| SPP1 | 0.175 | 0.506 | -0.0139 | 0.958 | 0.18 | 0.493 |
| C3 | -0.208 | 0.254 | 0.00268 | 0.988 | 0.142 | 0.433 |
| FN1 | 0.258 | 0.292 | 0.101 | 0.679 | 0.061 | 0.803 |
| COL6A3 | 0.218 | 0.628 | 0.0634 | 0.888 | -0.0385 | 0.932 |
| MBP | -0.299 | 0.326 | 0.0783 | 0.797 | -0.0795 | 0.794 |
| PLP1 | -0.261 | 0.367 | 0.00829 | 0.977 | -0.19 | 0.51 |
| ENPP6 | -0.179 | 0.515 | -0.0508 | 0.853 | -0.207 | 0.452 |

| | PC5 | | | | | |
|---|---|---|---|---|---|---|
| | Az | | Epo | | Az+Epo | |
| | Log2 fold change | P-value | Log2 fold change | P-value | Log2 fold change | P-value |
| SLC1A3 | 0.788 | 0.126 | 0.767 | 0.136 | 0.734 | 0.154 |
| PIK3R1 | 0.721 | 0.116 | 0.735 | 0.109 | 0.703 | 0.125 |
| CAMK2G | 0.58 | 0.201 | 0.708 | 0.12 | 0.666 | 0.143 |
| MMP2 | 0.568 | 0.189 | 0.553 | 0.2 | 0.646 | 0.135 |
| SOD1 | 0.691 | 0.139 | 0.703 | 0.132 | 0.596 | 0.201 |
| SERPINE2 | 0.566 | 0.233 | 0.728 | 0.126 | 0.583 | 0.219 |
| CLDN5 | 0.459 | 0.259 | 0.462 | 0.256 | 0.494 | 0.225 |
| ETV5 | 0.604 | 0.123 | 0.576 | 0.141 | 0.444 | 0.255 |
| FN1 | 0.258 | 0.292 | 0.101 | 0.679 | 0.061 | 0.803 |
| GAS6 | -0.0303 | 0.885 | 0.0649 | 0.758 | 0.0609 | 0.772 |
| ITGA7 | -0.142 | 0.422 | 0.0229 | 0.897 | 0.0515 | 0.771 |
| S100B | 0.0409 | 0.872 | 0.192 | 0.45 | 0.0233 | 0.927 |
| CXCL10 | 0.367 | 0.162 | 0.396 | 0.133 | -0.0229 | 0.93 |
| STAT1 | 0.165 | 0.503 | 0.174 | 0.479 | -0.0292 | 0.905 |
| CD86 | 0.0989 | 0.613 | 0.0881 | 0.652 | -0.0656 | 0.737 |

| | PC6 | | | | | |
|---|---|---|---|---|---|---|
| | Az | | Epo | | Az+Epo | |
| | Log2 fold change | P-value | Log2 fold change | P-value | Log2 fold change | P-value |
| CTSB | 0.797 | 0.193 | 0.863 | 0.159 | 0.832 | 0.175 |
| MMP14 | 0.87 | 0.0561 | 0.794 | 0.0809 | 0.795 | 0.0803 |
| TYROBP | 0.465 | 0.301 | 0.585 | 0.194 | 0.753 | 0.0954 |
| GLUL | 0.727 | 0.141 | 0.716 | 0.147 | 0.736 | 0.136 |
| LGALS3 | 0.788 | 0.134 | 0.695 | 0.186 | 0.713 | 0.175 |
| PRKAG1 | 0.615 | 0.102 | 0.578 | 0.124 | 0.594 | 0.114 |
| CLDN5 | 0.459 | 0.259 | 0.462 | 0.256 | 0.494 | 0.225 |
| SPP1 | 0.175 | 0.506 | -0.0139 | 0.958 | 0.18 | 0.493 |
| C3 | -0.208 | 0.254 | 0.00268 | 0.988 | 0.142 | 0.433 |
| GPR34 | 0.0855 | 0.69 | 0.233 | 0.279 | 0.0418 | 0.845 |
| PLA2G4A | 0.125 | 0.493 | 0.191 | 0.294 | 0.00424 | 0.981 |
| ITGAX | -0.0126 | 0.949 | 0.0651 | 0.739 | 0.00418 | 0.983 |
| PSMB9 | 0.0943 | 0.711 | 0.163 | 0.523 | -0.0177 | 0.945 |
| STAT1 | 0.165 | 0.503 | 0.174 | 0.479 | -0.0292 | 0.905 |
| CSF1R | -0.0127 | 0.957 | 0.117 | 0.622 | -0.0589 | 0.804 |

| | PC7 | | | | | |
|---|---|---|---|---|---|---|
| | Az | | Epo | | Az+Epo | |
| | **Log2 fold change** | **P-value** | **Log2 fold change** | **P-value** | **Log2 fold change** | **P-value** |
| CTSB | 0.797 | 0.193 | 0.863 | 0.159 | 0.832 | 0.175 |
| TYROBP | 0.465 | 0.301 | 0.585 | 0.194 | 0.753 | 0.0954 |
| ATP6V1B2 | 0.757 | 0.123 | 0.757 | 0.123 | 0.665 | 0.175 |
| SOX9 | 0.79 | 0.0882 | 0.638 | 0.167 | 0.628 | 0.174 |
| PRKAG1 | 0.615 | 0.102 | 0.578 | 0.124 | 0.594 | 0.114 |
| TSC1 | 0.656 | 0.0855 | 0.538 | 0.157 | 0.575 | 0.131 |
| ABL1 | 0.716 | 0.0859 | 0.606 | 0.145 | 0.529 | 0.202 |
| GSN | 0.71 | 0.101 | 0.565 | 0.191 | 0.468 | 0.278 |
| DLX1 | 0.823 | 0.0206 | 0.196 | 0.577 | 0.328 | 0.35 |
| CTSD | 0.0321 | 0.755 | 0.104 | 0.314 | 0.196 | 0.0594 |
| SPP1 | 0.175 | 0.506 | -0.0139 | 0.958 | 0.18 | 0.493 |
| C3 | -0.208 | 0.254 | 0.00268 | 0.988 | 0.142 | 0.433 |
| DLX2 | 0.514 | 0.0958 | 0.0192 | 0.95 | 0.00727 | 0.981 |
| CXCL10 | 0.367 | 0.162 | 0.396 | 0.133 | -0.0229 | 0.93 |
| SOX2 | 0.234 | 0.275 | 0.0611 | 0.775 | -0.0572 | 0.789 |

**Table S4. Differentially expressed genes between OGD and treatments globally, and in the white matter, deep grey matter, and cortex.** Genes were considered differentially expressed when p-value <0.05.

**Global**

**OGD vs. Control**

| Genes | Log2 fold change | P-value |
|---|---|---|
| MAFB | 4.63 | 1.5E-15 |
| MAG | 3.28 | 4.92E-10 |
| CCL5 | 1.4 | 1.17E-06 |
| AHCYL1 | 1.1 | 1.77E-06 |
| MARCO | 0.99 | 0.000105 |
| CDKN1A | 0.732 | 2.69E-06 |
| S100B | 0.6 | 0.0201 |
| ABCG2 | 0.587 | 2.41E-08 |
| ITGAX | 0.575 | 0.00417 |
| CD86 | 0.55 | 0.00589 |
| CASP8 | 0.469 | 0.0144 |
| MMP19 | 0.456 | 0.00117 |
| MERTK | 0.421 | 0.0447 |
| TSPO | 0.397 | 0.0127 |
| TFEC | 0.375 | 0.0473 |
| GUSB | 0.272 | 0.0104 |
| CTSD | 0.228 | 0.0294 |
| PARP1 | 0.196 | 0.000672 |
| UBE4B | 0.149 | 0.0051 |
| ATP6V1A | -0.182 | 0.00476 |
| ULK1 | -0.305 | 0.017 |
| MMP16 | -0.391 | 0.011 |
| GPR84 | -0.395 | 0.0097 |
| TENM4 | -0.404 | 0.00702 |
| NCAM2 | -0.438 | 0.0253 |
| IGF1 | -0.459 | 0.0112 |
| NEFH | -0.466 | 0.000188 |
| SLA | -0.494 | 0.019 |
| SPTAN1 | -0.512 | 0.00702 |
| NRG1 | -0.54 | 0.00841 |
| GNG2 | -0.56 | 1.64E-06 |
| RORB | -0.585 | 0.000216 |
| PRKCZ | -0.611 | 0.00696 |
| MEF2A | -0.629 | 0.00389 |
| CDS1 | -0.631 | 1.75E-06 |
| THY1 | -0.639 | 0.0108 |
| MEAF6 | -0.641 | 0.000439 |
| SYT4 | -0.693 | 0.0337 |
| CUX2 | -0.694 | 0.0436 |
| VEGFA | -0.71 | 9.61E-05 |
| EGR1 | -0.713 | 0.00256 |
| MAP2 | -0.716 | 4.6E-08 |
| ZNF365 | -0.723 | 0.000342 |
| C3 | -0.742 | 9.39E-05 |
| KCNJ10 | -0.754 | 0.0393 |
| MMP24 | -0.776 | 0.000989 |
| CNTNAP1 | -0.778 | 0.0002 |
| MAPK10 | -0.788 | 0.000464 |
| STMN1 | -0.802 | 0.00133 |
| RTN1 | -0.804 | 0.00261 |
| FGF13 | -0.819 | 0.00263 |
| RELN | -0.821 | 0.00677 |
| OPCML | -0.833 | 0.000149 |
| LRRC7 | -0.849 | 0.000628 |
| PTEN | -0.854 | 0.0388 |
| MAPK9 | -0.865 | 0.0318 |
| OLFM3 | -0.868 | 0.000235 |
| SH3GL2 | -0.885 | 0.000228 |
| CAMK2B | -0.889 | 0.00209 |
| NAP1L2 | -0.893 | 5.71E-08 |
| STMN4 | -0.92 | 0.000841 |
| PPM1L | -0.925 | 0.0235 |
| CDK5R1 | -0.933 | 0.000266 |
| SOD2 | -0.949 | 0.0249 |
| UCHL1 | -0.958 | 1.94E-07 |
| STXBP1 | -0.97 | 7.63E-06 |
| TBR1 | -0.981 | 0.000726 |
| TRIM46 | -0.982 | 0.000149 |
| ACSL4 | -0.998 | 0.0197 |
| XK | -0.999 | 3.32E-08 |
| DCLK1 | -1.01 | 0.0307 |
| XBP1 | -1.02 | 0.0137 |
| YWHAH | -1.03 | 2.86E-05 |
| SHANK2 | -1.04 | 7.62E-05 |
| NEFL | -1.1 | 2.41E-06 |
| SYT1 | -1.1 | 0.000242 |
| PSEN2 | -1.13 | 0.00228 |
| NPTN | -1.14 | 0.0182 |
| NTF3 | -1.15 | 0.000436 |
| SCN2A | -1.21 | 4.13E-05 |
| YWHAZ | -1.23 | 0.0243 |
| SNCA | -1.26 | 5.12E-06 |
| SLC12A5 | -1.27 | 1.33E-05 |
| SYN1 | -1.29 | 1.04E-05 |
| YWHAG | -1.29 | 0.0131 |
| SYNGR1 | -1.36 | 9.11E-07 |
| CD74 | -1.39 | 3.23E-12 |
| CAMK4 | -1.44 | 3.95E-07 |
| EGR2 | -1.55 | 1.15E-06 |
| BDNF | -1.55 | 0.000002 |
| LINGO1 | -1.56 | 3.77E-06 |
| RASGRF1 | -1.6 | 7.09E-09 |

**Az vs. OGD**

| Genes | Log2 fold change | P-value |
|---|---|---|
| DLX1 | 0.823 | 0.0206 |
| CCL2 | 0.702 | 0.0123 |
| MMP12 | 0.629 | 0.029 |
| GPR84 | 0.409 | 0.00757 |
| FLT1 | 0.343 | 0.0475 |
| ABCG2 | -0.192 | 0.047 |
| CASP3 | -0.509 | 0.0175 |
| CDKN1A | -0.576 | 0.000155 |
| ISLR2 | -0.762 | 0.00664 |
| AHCYL1 | -1.04 | 5.9E-06 |
| MAG | -3.6 | 2.01E-11 |
| MAFB | -4.5 | 5.17E-15 |

**Epo vs. OGD**

| Genes | Log2 fold change | P-value |
|---|---|---|
| CDKN1A | -0.508 | 0.000751 |
| CASP3 | -0.67 | 0.00199 |
| ISLR2 | -0.848 | 0.00264 |
| AHCYL1 | -0.882 | 9.13E-05 |
| MAG | -3.27 | 5.06E-10 |
| MAFB | -4.49 | 5.94E-15 |

**Az+Epo vs. OGD**

| Genes | Log2 fold change | P-value |
|---|---|---|
| SRPRA | 0.758 | 0.045 |
| GPR84 | 0.489 | 0.00149 |
| CD74 | 0.459 | 0.00897 |
| ADGRG1 | 0.24 | 0.023 |
| TIMP2 | 0.19 | 0.046 |
| UBE4B | -0.149 | 0.00519 |
| ABCG2 | -0.244 | 0.0123 |
| CDKN1A | -0.436 | 0.00355 |
| ISLR2 | -0.587 | 0.0348 |
| CASP3 | -0.61 | 0.00463 |
| AHCYL1 | -0.924 | 4.48E-05 |
| MAG | -3.49 | 6.18E-11 |
| MAFB | -4.37 | 1.9E-14 |

**White Matter**

**OGD vs. Control**

| Genes | Log2 fold change | P-value |
|---|---|---|
| CCL5 | 1.4 | 1.49E-05 |
| COL6A3 | 1.32 | 0.0461 |
| MARCO | 1.26 | 0.000368 |
| S100B | 1.08 | 2.09E-05 |
| ITGAX | 0.613 | 6.63E-05 |
| TNF | 0.551 | 0.00426 |
| CD86 | 0.535 | 0.000109 |
| MMP19 | 0.524 | 0.00651 |
| FN1 | 0.519 | 0.0163 |
| GSN | 0.5 | 0.000425 |
| CLDN5 | 0.5 | 0.00058 |
| CASP8 | 0.499 | 0.00162 |
| ITGA5 | 0.485 | 9.84E-05 |
| PLP1 | 0.478 | 0.00611 |
| PLXNB3 | 0.478 | 0.00627 |
| EGFR | 0.477 | 1.17E-05 |
| TSPO | 0.471 | 0.000192 |
| GAS6 | 0.47 | 0.0018 |
| SOX10 | 0.461 | 0.0128 |
| JAM3 | 0.459 | 0.000626 |
| FTH1 | 0.434 | 2.02E-05 |
| MMP2 | 0.424 | 0.00551 |
| SOX2 | 0.421 | 0.000799 |
| PLXNC1 | 0.415 | 0.00183 |
| MERTK | 0.403 | 0.00177 |
| LGALS3 | 0.402 | 0.0302 |
| ITGB5 | 0.399 | 0.000125 |
| IRF8 | 0.394 | 0.00281 |
| SOX9 | 0.393 | 0.00155 |
| CNTF | 0.39 | 0.00226 |
| ABCG2 | 0.389 | 0.00195 |
| IL6R | 0.389 | 0.0308 |
| AXL | 0.388 | 0.00185 |
| CNP | 0.381 | 0.0319 |
| UGT8 | 0.373 | 0.0334 |
| LMNA | 0.372 | 0.00088 |
| TFEC | 0.37 | 0.0125 |
| CYBB | 0.368 | 0.00133 |
| ITGA7 | 0.366 | 0.000408 |
| PLEKHM1 | 0.346 | 2.56E-05 |
| MYRF | 0.339 | 0.0314 |
| SPI1 | 0.33 | 0.0384 |
| NLRP3 | 0.33 | 0.0465 |
| MMP14 | 0.328 | 0.000959 |
| SERPINI1 | 0.325 | 0.0393 |
| NAGLU | 0.323 | 0.0199 |
| GAA | 0.322 | 0.00254 |
| TREM2 | 0.318 | 0.0204 |
| CTSB | 0.31 | 0.0294 |
| CDKN1A | 0.309 | 0.0245 |
| SIRT2 | 0.308 | 0.0141 |
| PMP22 | 0.298 | 0.0014 |
| GBA | 0.291 | 0.00567 |
| C5AR1 | 0.288 | 0.00751 |
| CH25H | 0.283 | 0.0331 |
| MAPK3 | 0.28 | 0.00481 |
| TLR2 | 0.276 | 0.00831 |
| ABL1 | 0.272 | 0.000824 |
| AHCYL1 | 0.267 | 0.00326 |
| ACADSB | 0.259 | 0.016 |
| SCAMP2 | 0.253 | 0.0154 |
| RHOA | 0.25 | 0.00405 |
| PSEN1 | 0.239 | 0.00169 |
| TLR4 | 0.237 | 0.0128 |
| IKBKB | 0.236 | 0.00543 |
| CTSD | 0.229 | 0.0447 |
| MAFB | 0.223 | 0.0388 |
| CASP2 | 0.211 | 0.0457 |
| PRKAB1 | 0.197 | 0.00792 |
| ARHGEF10 | 0.185 | 0.0118 |
| CD9 | 0.171 | 0.0255 |
| TGFBR1 | 0.171 | 0.0467 |
| PARP1 | 0.129 | 0.0182 |
| PARP2 | 0.127 | 0.0507 |
| TIMP2 | 0.121 | 0.0307 |
| PTEN | -0.171 | 0.00104 |
| ATP6V1A | -0.179 | 0.0481 |
| DMD | -0.224 | 0.0518 |
| SOD2 | -0.318 | 6.17E-06 |
| MAPK9 | -0.334 | 0.00495 |
| VEGFA | -0.348 | 0.019 |
| MMP16 | -0.36 | 0.00644 |
| SPTAN1 | -0.361 | 0.00027 |
| XBP1 | -0.366 | 9.44E-06 |
| PPM1L | -0.38 | 5.04E-05 |
| DCLK1 | -0.382 | 0.00199 |
| IGF1 | -0.407 | 0.0219 |
| NPTN | -0.424 | 6.56E-05 |
| ACSL4 | -0.445 | 0.00162 |
| ZNF365 | -0.445 | 0.00219 |
| NCAM2 | -0.452 | 0.0222 |
| CDS1 | -0.465 | 0.00021 |
| MEF2A | -0.466 | 0.00386 |
| CUX2 | -0.467 | 0.045 |
| YWHAZ | -0.484 | 0.000293 |
| CASP3 | -0.487 | 0.000336 |
| NEFH | -0.5 | 3.25E-06 |
| RORB | -0.523 | 0.00047 |
| PRKCZ | -0.534 | 0.000289 |
| YWHAG | -0.545 | 1.97E-06 |
| EGR1 | -0.564 | 0.00568 |
| MEAF6 | -0.589 | 0.000385 |
| THY1 | -0.61 | 0.000517 |
| MAP2 | -0.615 | 6.89E-05 |
| CAMK2B | -0.618 | 0.00021 |
| PSEN2 | -0.62 | 1.13E-06 |
| MMP24 | -0.646 | 0.00018 |
| LRRC7 | -0.705 | 0.000114 |
| CNTNAP1 | -0.71 | 2.74E-05 |
| GNG2 | -0.746 | 7.46E-06 |
| SYT4 | -0.748 | 0.0284 |
| FGF13 | -0.754 | 0.000241 |
| RELN | -0.761 | 0.00457 |
| STMN1 | -0.765 | 3.68E-05 |
| MAPK10 | -0.766 | 3.31E-05 |
| RTN1 | -0.766 | 0.000143 |
| OPCML | -0.775 | 7.39E-05 |
| OLFM3 | -0.802 | 0.00375 |
| NAP1L2 | -0.803 | 4.33E-08 |
| C3 | -0.834 | 0.00495 |
| TRIM46 | -0.86 | 4.84E-05 |
| STMN4 | -0.869 | 2.45E-05 |
| UCHL1 | -0.878 | 7.05E-06 |
| CDK5R1 | -0.885 | 6.64E-05 |
| NTF3 | -0.9 | 0.00779 |
| SH3GL2 | -0.952 | 7.72E-05 |
| STXBP1 | -0.958 | 2.89E-05 |
| ISLR2 | -0.997 | 0.000178 |
| SHANK2 | -0.998 | 1.13E-05 |
| YWHAH | -1.01 | 1.14E-06 |
| XK | -1.03 | 4.05E-06 |
| SYNGR1 | -1.05 | 5.11E-07 |
| CD74 | -1.14 | 5.38E-05 |
| SYT1 | -1.16 | 2.67E-05 |
| TBR1 | -1.18 | 0.000161 |
| SCN2A | -1.2 | 1.03E-05 |
| SYN1 | -1.25 | 8.39E-06 |
| NEFL | -1.27 | 4.32E-05 |
| LINGO1 | -1.28 | 3.6E-06 |
| SLC12A5 | -1.33 | 3.01E-05 |
| EGR2 | -1.35 | 5.08E-05 |
| RASGRF1 | -1.5 | 7.99E-07 |
| SNCA | -1.51 | 9.12E-06 |
| CAMK4 | -1.61 | 4.76E-06 |
| BDNF | -2.11 | 4.17E-05 |

**Az vs. OGD**

| Genes | Log2 fold change | P-value |
|---|---|---|
| PPM1L | 0.239 | 0.00502 |
| PARP1 | 0.174 | 0.00218 |
| PSEN1 | 0.17 | 0.0193 |
| SOD2 | 0.165 | 0.00679 |
| ITGA7 | -0.196 | 0.0379 |
| CLDN5 | -0.298 | 0.0273 |
| CNP | -0.353 | 0.0452 |
| NGFR | -0.451 | 0.0369 |
| RELN | -0.673 | 0.0112 |

**Epo vs. OGD**

| Genes | Log2 fold change | P-value |
|---|---|---|
| ARHGEF10 | -0.176 | 0.0163 |
| SERPINI1 | -0.464 | 0.0047 |
| RELN | -0.548 | 0.0346 |
| NTF3 | -0.763 | 0.0224 |
| COL6A3 | -1.31 | 0.047 |

**Az+Epo vs. OGD**

| Genes | Log2 fold change | P-value |
|---|---|---|
| CD74 | 0.54 | 0.031 |
| GPR84 | 0.433 | 0.0116 |
| ADGRG1 | 0.348 | 0.0227 |
| CD14 | 0.275 | 0.0377 |
| TSPO | 0.254 | 0.0256 |
| PSEN2 | 0.246 | 0.0184 |
| SRPRA | 0.232 | 0.0222 |
| CTSD | 0.223 | 0.0498 |
| CASP9 | 0.202 | 0.0375 |
| TIMP2 | 0.126 | 0.0254 |
| ETV5 | -0.195 | 0.0347 |
| HDAC2 | -0.211 | 0.0377 |
| JAM3 | -0.243 | 0.0488 |
| EGFR | -0.269 | 0.00506 |
| HGF | -0.278 | 0.0426 |
| MARCO | -0.734 | 0.0233 |

#### Deep Grey Matter

**OGD vs. Control**

| Genes | Log2 fold change | P-value |
|---|---|---|
| IL10 | 0.556 | 0.00583 |
| ITGAX | 0.485 | 0.00208 |
| CD86 | 0.431 | 0.00458 |
| PLXNC1 | 0.427 | 0.00228 |
| CDKN1A | 0.416 | 0.0051 |
| ITGA7 | 0.395 | 0.0121 |
| JAM3 | 0.372 | 0.0116 |
| GAS6 | 0.372 | 0.027 |
| EGFR | 0.368 | 0.00671 |
| CASP8 | 0.368 | 0.0234 |
| ABCG2 | 0.341 | 0.00092 |
| MERTK | 0.336 | 0.0242 |
| GSN | 0.336 | 0.0351 |
| CTSB | 0.335 | 0.0279 |
| MMP2 | 0.325 | 0.0434 |
| CNTF | 0.316 | 0.0136 |
| SPI1 | 0.314 | 0.0287 |
| FTH1 | 0.311 | 0.00141 |
| CLDN5 | 0.302 | 0.0245 |
| GAA | 0.3 | 0.0168 |
| SLC1A3 | 0.3 | 0.0271 |
| ACADSB | 0.295 | 0.00855 |
| FIG4 | 0.283 | 0.0456 |
| PLEKHM1 | 0.276 | 0.00156 |
| PSEN1 | 0.274 | 0.00297 |
| NLRP3 | 0.272 | 0.0481 |
| ITGB5 | 0.264 | 0.0159 |
| GBA | 0.262 | 0.0273 |
| MAPK3 | 0.248 | 0.0123 |
| C5AR1 | 0.239 | 0.0492 |
| GNPTAB | 0.236 | 0.0179 |
| LMNA | 0.235 | 0.039 |
| BCL2L11 | 0.21 | 0.0334 |
| PTEN | -0.121 | 0.0387 |
| SOD2 | -0.188 | 0.0158 |
| CD40 | -0.225 | 0.0202 |
| NPTN | -0.255 | 0.0481 |
| CDS1 | -0.27 | 0.0196 |
| TENM4 | -0.274 | 0.0346 |
| CD14 | -0.307 | 0.0369 |
| GNG2 | -0.311 | 0.0389 |
| YWHAG | -0.317 | 0.0134 |
| XBP1 | -0.332 | 9.56E-05 |
| MEAF6 | -0.344 | 0.0283 |
| ZNF365 | -0.363 | 0.0414 |
| MEF2A | -0.372 | 0.049 |
| IGF1 | -0.395 | 0.0368 |
| CASP3 | -0.414 | 0.0319 |
| PSEN2 | -0.424 | 0.000533 |
| MAP2 | -0.448 | 0.0227 |
| STMN1 | -0.471 | 0.0486 |
| MAPK10 | -0.476 | 0.0216 |
| NAP1L2 | -0.493 | 0.000103 |
| GPR84 | -0.545 | 0.0206 |
| VEGFA | -0.546 | 0.0212 |
| UCHL1 | -0.556 | 0.0152 |
| RTN1 | -0.566 | 0.0424 |
| CDK5R1 | -0.596 | 0.0173 |
| STXBP1 | -0.626 | 0.0139 |
| YWHAH | -0.65 | 0.0005 |
| XK | -0.68 | 0.00023 |
| NEFL | -0.681 | 0.00593 |
| SHANK2 | -0.704 | 0.0143 |
| EGR1 | -0.71 | 0.00753 |
| SNCA | -0.766 | 0.0041 |
| OLFM3 | -0.79 | 0.00159 |
| SYNGR1 | -0.807 | 0.0137 |
| SYT1 | -0.828 | 0.00791 |
| C3 | -0.838 | 0.00482 |
| SCN2A | -0.862 | 0.00907 |
| SYN1 | -0.91 | 0.00415 |
| SLC12A5 | -0.997 | 0.000829 |
| LINGO1 | -1.09 | 0.0107 |
| CAMK4 | -1.1 | 0.000306 |
| RASGRF1 | -1.26 | 9.19E-05 |
| CD74 | -1.48 | 1.18E-06 |
| EGR2 | -1.64 | 0.000224 |

**Az vs. OGD**

| Genes | Log2 fold change | P-value |
|---|---|---|
| CXCL10 | 0.739 | 0.0201 |
| SOX2 | 0.269 | 0.0485 |
| CD40 | 0.207 | 0.0313 |
| PARP1 | 0.181 | 0.00614 |
| NF1 | 0.179 | 0.0445 |
| MTOR | -0.196 | 0.0401 |
| C5AR1 | -0.259 | 0.0338 |
| CDKN1A | -0.303 | 0.0339 |
| TYROBP | -0.331 | 0.0243 |
| TREM1 | -0.447 | 0.0368 |
| CNP | -0.571 | 0.014 |
| NGFR | -0.632 | 0.00478 |

**Epo vs. OGD**

| Genes | Log2 fold change | P-value |
|---|---|---|
| CXCL10 | 0.827 | 0.0102 |
| MTOR | -0.204 | 0.0331 |
| NAGLU | -0.326 | 0.023 |
| NGFR | -0.491 | 0.0227 |

**Az+Epo vs. OGD**

| Genes | Log2 fold change | P-value |
|---|---|---|
| MAPK14 | -0.103 | 0.0467 |
| BCL2L11 | -0.225 | 0.0232 |
| EGFR | -0.257 | 0.0494 |
| ADRA2A | -0.533 | 0.0482 |

#### Cortex

**OGD vs. Control**

| Genes | Log2 fold change | P-value |
|---|---|---|
| MAFB | 6.29 | 1.71E-07 |
| CCL7 | 2.69 | 0.000451 |
| AHCYL1 | 2.19 | 0.000103 |
| MARCO | 1.39 | 0.0096 |
| CDKN1A | 1.29 | 0.000586 |
| ABCG2 | 1.03 | 8.19E-05 |
| GUSB | 0.35 | 0.00546 |
| PARP1 | 0.343 | 0.0156 |
| UBE4B | 0.314 | 0.00866 |
| ATP6V1A | -0.311 | 0.00307 |
| GNG2 | -0.577 | 0.00224 |
| RORB | -0.772 | 0.000232 |
| OLFM3 | -0.969 | 0.0447 |
| MAP2 | -0.987 | 3.28E-06 |
| STXBP1 | -1.17 | 0.000225 |
| XK | -1.23 | 0.00332 |
| DLX1 | -1.24 | 0.00315 |
| UCHL1 | -1.28 | 7.42E-05 |
| BDNF | -1.51 | 0.0254 |
| CD74 | -1.54 | 0.00044 |
| EGR2 | -1.62 | 0.00164 |
| NEFH | -1.74 | 0.035 |
| DLX2 | -1.9 | 0.0496 |
| TENM4 | -1.96 | 0.0472 |
| MMP16 | -2.03 | 0.0468 |
| CDS1 | -2.04 | 0.0139 |
| CUX2 | -2.23 | 0.0396 |
| PSEN2 | -2.33 | 0.0358 |
| TBR1 | -2.36 | 0.0191 |
| NRG1 | -2.36 | 0.0267 |
| MEAF6 | -2.39 | 0.036 |
| RELN | -2.41 | 0.032 |
| ADRA2A | -2.45 | 0.0068 |
| VEGFA | -2.45 | 0.0235 |
| NAP1L2 | -2.5 | 0.00973 |
| SLA | -2.5 | 0.0335 |
| SPTAN1 | -2.5 | 0.0471 |
| NTF3 | -2.59 | 0.00668 |
| MMP24 | -2.59 | 0.031 |
| CNTNAP1 | -2.6 | 0.0203 |
| ZNF365 | -2.61 | 0.0225 |
| SHANK2 | -2.64 | 0.0199 |
| TRIM46 | -2.65 | 0.0192 |
| MAPK10 | -2.65 | 0.0376 |
| SH3GL2 | -2.66 | 0.0258 |
| MEF2A | -2.72 | 0.0448 |
| OPCML | -2.74 | 0.0258 |
| FGF13 | -2.77 | 0.0397 |
| NEFL | -2.81 | 0.0197 |
| CAMK2B | -2.81 | 0.0275 |
| SLC12A5 | -2.95 | 0.0155 |
| SNCA | -2.95 | 0.0189 |
| LRRC7 | -2.95 | 0.03 |
| CDK5R1 | -2.95 | 0.0322 |
| RASGRF1 | -2.96 | 0.00311 |
| SCN2A | -3.01 | 0.0182 |
| SYT1 | -3.08 | 0.0312 |
| STMN4 | -3.13 | 0.0372 |
| CAMK4 | -3.14 | 0.0137 |
| SYN1 | -3.16 | 0.015 |
| YWHAH | -3.17 | 0.0293 |
| STMN1 | -3.19 | 0.0462 |
| SYNGR1 | -3.25 | 0.00692 |
| LINGO1 | -3.35 | 0.00685 |

**Az vs. OGD**

| Genes | Log2 fold change | P-value |
|---|---|---|
| MMP12 | 1.33 | 0.0298 |
| NPAS4 | 1.11 | 0.0374 |
| GPR84 | 0.835 | 0.00845 |
| RORB | 0.484 | 0.0126 |
| ABCG2 | -0.508 | 0.0283 |
| CASP3 | -1.06 | 0.00995 |
| CDKN1A | -1.21 | 0.00112 |
| ISLR2 | -1.31 | 0.00356 |
| CCR2 | -2 | 0.00593 |
| AHCYL1 | -2.15 | 0.000132 |
| MAFB | -6.15 | 2.54E-07 |

**Epo vs. OGD**

| Genes | Log2 fold change | P-value |
|---|---|---|
| NPAS4 | 1.18 | 0.028 |
| DLX1 | 0.832 | 0.0386 |
| ABCG2 | -0.484 | 0.0359 |
| CDKN1A | -1.13 | 0.00214 |
| CASP3 | -1.16 | 0.00513 |
| ISLR2 | -1.28 | 0.00446 |
| CCR2 | -1.8 | 0.012 |
| AHCYL1 | -2.03 | 0.000249 |
| MAFB | -6.18 | 2.32E-07 |

**Az+Epo vs. OGD**

| Genes | Log2 fold change | P-value |
|---|---|---|
| GPR84 | 0.869 | 0.00639 |
| ABCG2 | -0.596 | 0.0114 |
| ISLR2 | -0.992 | 0.0225 |
| CASP3 | -1.03 | 0.0118 |
| CDKN1A | -1.12 | 0.00224 |
| AHCYL1 | -2.1 | 0.000173 |
| CCR2 | -2.25 | 0.00238 |
| MAFB | -6.05 | 3.26E-07 |
